# Telomerase-independent maintenance of telomere length in a vertebrate

**DOI:** 10.1101/2022.03.25.485759

**Authors:** Qinghao Yu, Phillip B. Gates, Samuel Rogers, Ivan Mikicic, Ahmed Elewa, Florian Salomon, Martina Lachnit, Antonio Caldarelli, Neftali Flores-Rodriguez, Anthony J Cesare, Andras Simon, Maximina H. Yun

**Affiliations:** Technische Universität Dresden, CRTD/Center for Regenerative Therapies Dresden; Dresden, Germany; Institute of Structural and Molecular Biology, University College London; UK; Genome Integrity Unit, Children’s Medical Research Institute, University of Sydney; Sydney, New South Wales, Australia; Department of Cell and Molecular Biology, Karolinska Institute; Stockholm, Sweden; Australian Centre for Microscopy and Microanalysis, University of Sydney, Sydney, New South Wales, Australia; Max Planck Institute for Molecular Cell Biology and Genetics; Dresden, Germany

## Abstract

Telomere shortening places a key limitation on cell proliferation^1^. In all vertebrates explored to date, this limitation is overcome by telomerase-dependent telomere extension. Failure to maintain telomere length results in premature ageing and functional impairments in highly replicative cell populations as telomeres erode^2^. Alternative lengthening of telomeres (ALT), a telomerase-independent mechanism, compensates for telomere loss in a subset of human cancer cell lines ^2^. Here, we demonstrate that the highly regenerative newt *Pleurodeles waltl* lacks telomerase activity, contains telomeres distinct from all known vertebrates in both sequence and structure, and deploys ALT for physiological telomere maintenance. This constitutes the first report of telomerase-independent resolution of the end-replication problem at the whole-organism level within *Chordata*.

**One-Sentence Summary:** *P. waltl* telomere biology is distinct amongst vertebrates and uses ALT at the whole-organism level.

## Main Text

Vertebrate telomeres consist of serial 5’-TTAGGG-3’ repeats located at chromosome ends(*1*). Due to the semiconservative nature of DNA replication, telomeres shorten with each cell division(*2*). This ultimately leads to telomere erosion, activation of the DNA damage response and triggering of replicative senescence, processes associated with ageing(*3*). While normal somatic cells cannot overcome this problem, highly replicative populations including stem, germ and immortalised cells maintain telomere length through the reverse-transcriptase telomerase(*4*). In the absence of telomerase, a subset of cancer cells adopt an alternative lengthening of telomeres (ALT) mechanism which relies on homologous recombination(*5*). However, ALT has never been reported to operate at the whole-organism level(*6*).

Telomere length maintenance poses a significant challenge for adult regeneration(*7*). In organisms with unlimited regenerative capacity such as planarian flatworms, this is overcome through telomerase upregulation(*8*). Within vertebrates, salamanders exhibit indefinite regenerative abilities(*9*), though the mechanism of telomere length maintenance in these animals remains unknown. Of note, salamander regenerative tissue is highly resistant to tumor formation despite exhibiting several hallmarks of cancer(*10*), and cells derived from the limb blastema, the structure that gives rise to the regenerated extremity, do not exhibit signs of replicative senescence(*9*). We therefore explored how salamanders solve the end-replication problem.

First, we surveyed telomerase activity using TRAP assays (fig.S1) in developing (Fig. 1, A and B) and regenerating limb tissues (Fig. 1, C and D) of two related salamander species, the axolotl *Ambystoma mexicanum* and the newt *Pleurodeles waltl. A. mexicanum* exhibits clear telomerase activity in larval extracts (Fig. 1A), limb epidermis and regenerating limbs (Fig. 1C), whereas telomerase activity is absent in corresponding *P. waltl* tissues (Fig. 1B and D). This cannot be attributed to the presence of an inhibitor in newt tissues, as combinations of axolotl and newt extracts do not impair axolotl telomerase activity (Fig. 1D). Thus, we asked whether the observed differences in telomerase activity could result from differences at the expression level. Mirroring its activity, axolotl *Tert* transcription is upregulated in larva, skin and regenerating limbs (fig. S2). By mining the axolotl genome and transcriptome(*11*) we identified the locus corresponding to the catalytic subunit of telomerase, *Tert*, and its flanking genes (Fig. 1E). Notably, homology-based blast iterations were not able to detect *Tert* in the *P. waltl* genome nor in transcriptomes derived from developing and regenerating newts, (*12*) despite identifying all other genes flanking the *Tert* locus (Fig. 1E), suggesting that *P. waltl Tert* has diverged beyond detection or was altogether lost. In agreement, *P. waltl Tert* is not detected by genomic DNA hybridisation against a highly conserved *Tert* region (fig. S3). Together, these results suggest that *A. mexicanum* overcomes telomere maintenance demands during development, homeostasis and regeneration by upregulating telomerase, and that a different mechanism is employed by *P. waltl*.

**Fig. 1.**
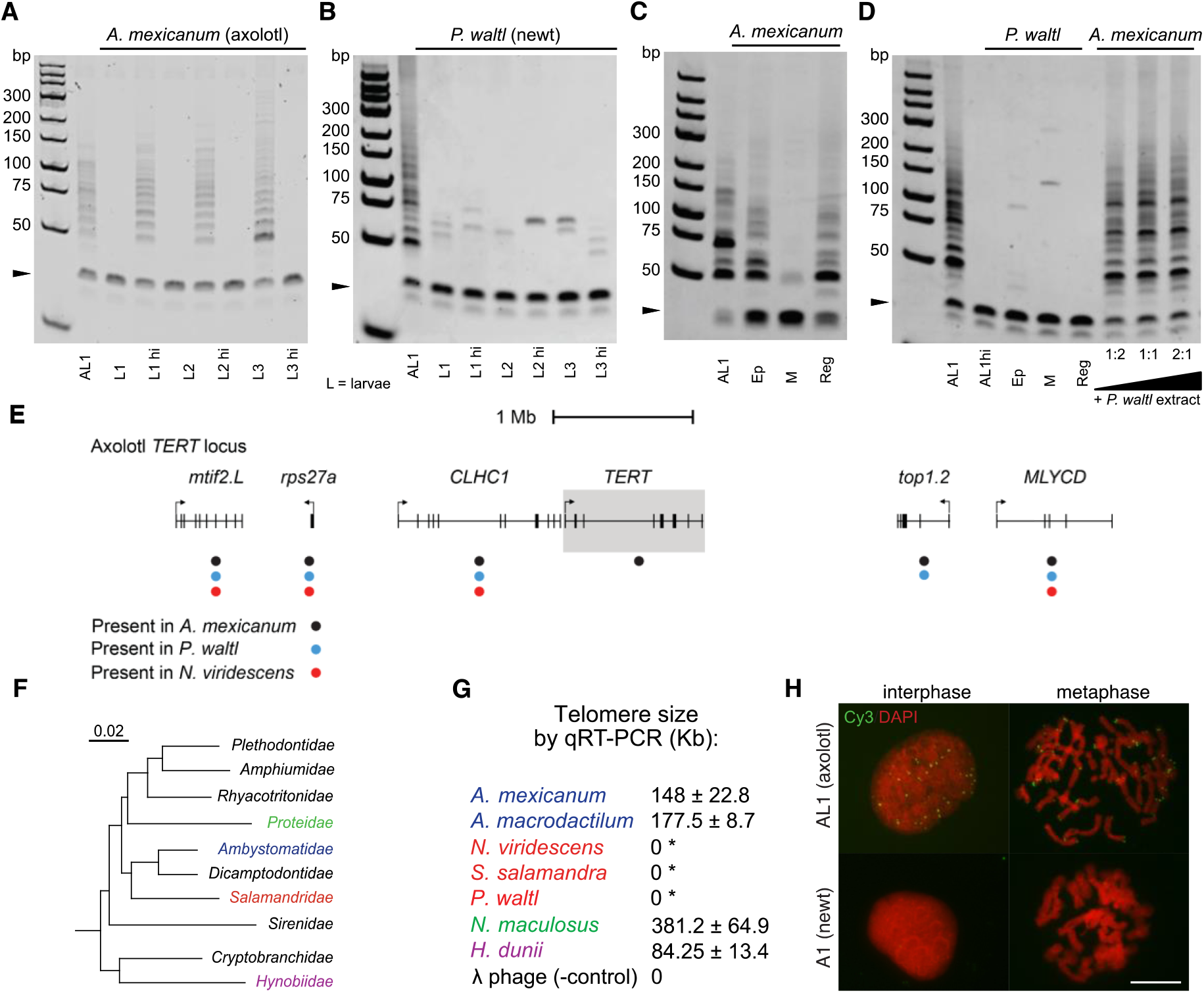
Absence of telomerase and canonical telomeres in *P. waltl*. (**A-D**) Telomerase enzymatic activity in *A. mexicanum* (**A, C, D**) and *P. waltl* (**B, D**) extracts from whole larva (A, B), from paired-matched intact limb skin (Ep), limb mesenchyme (M) and blastema (Reg) (C, D), and from *A. mexicanum* AL1 cells combined with increasing amounts of *P. waltl* skin extract (D) (n=6). A 36-bp amplification control is included in each reaction (arrowhead). AL1 cell extract is included in each experiment as positive control. hi=heat inactivation. (**E**) *P. waltl* lacks the *Tert* locus. Structure of *A. mexicanum Tert* locus and conserved flanking genes. The presence of each gene in *P. waltl* (genome and transcriptome) and *N. viridescens* (transcriptome) is shown (coloured dots). (**F**) Phylogenetic tree showing the placement of 10 salamander families, adapted from(*30*). For reference, the divergence between *Ambystomatidae* and *Dicamptodontidae* is estimated to have occurred 99.6 million years ago. The rate of mutation per nucleotide corresponding to 10 million years is indicated. (**G**) Relative telomeric DNA content calculated by genomic qPCR using telomere consensus and single-copy gene normaliser primers for the indicated species, representing various salamander families (coloured). All species belonging to *Salamandridae* (*) give qRT-PCR traces equivalent to negative control (λ phage) when telomere consensus primers are used. Values represent mean ± SD of ≥3 independent experiments. (**H**) FISH assay using a telomere consensus probe on metaphase spreads derived from axolotl AL1 and newt (*N. viridescens*) A1 cells (n=5).

We attempted to determine the absolute telomere size for both species as well as five close evolutionary relatives (Fig. 1F) using qPCR of the canonical TTAGGG vertebrate telomere sequence (Fig. 1G). This revealed that members of the *Ambystomatidae, Proteidae*, and *Hynobiidae* salamander families possess telomeres of canonical sequence that are 10-30 times longer than typical human telomeres. Detection of consensus telomere signal in *Hynobiidae* suggests that the ancestor of the modern crown-group salamanders possessed canonical vertebrate telomeres. Conversely, we failed to detect canonical telomere signal by qPCR in all three members of *Salamandridae*, the family comprising newts and true salamanders, including *P. waltl* (Fig. 1G). In agreement, Fluorescence *In Situ* Hybridization (FISH) of axolotl samples using a consensus telomere probe revealed punctate foci in spleen nuclei and at chromosome termini in mitotic spreads (Fig. 1H and fig. S4). In contrast, equivalent canonical telomere signals were absent in newt (*Notophthalmus viridescens*) cells. This was not due to a technical limitation as FISH signals against other newt repetitive elements were readily visualised (fig. S5). This is consistent with reports(*13*) demonstrating a lack of consensus telomere signal in another member of *Salamandridae, Cynops pyrrhogaster*, and with the failure to detect *Tert* in the *N. viridescens* transcriptome (Fig. 1E). Together, these results indicate that *Salamandridae* chromosome termini differ from other known vertebrates.

Next, we sought to determine the identity of *A. mexicanum* and *P. waltl* telomeres. Mining of PacBio long read-derived whole-genome sequences for *A. mexicanum*(*11*) and *P. waltl* (supplementary material) allowed us to retrieve the telomeric sequences for each species (Fig. 2A, fig. S6 and supplementary text). Critically, whereas axolotl telomeres consist of long tracks of TTAGGG vertebrate consensus hexamers (>90%, Fig. 2, A and B, fig. S7), *P. waltl* telomeres are heavily interspersed with variant telomere repeats (>50%, Fig. 2, A and C, fig. S7), a signature of ALT-mediated telomere maintenance(*14*). Notably, the ratio of variant versus canonical repeats in *P. waltl* telomeres is considerably higher than in any ALT^+^ cell reported thus far(*15, 16*). As in mammals(*17*), subtelomeric regions in both *A. mexicanum* and *P. waltl* contain a proportion of telomere variant repeats (Fig. 2, B and C, fig. S7), although this is elevated in *P. waltl*.

**Fig. 2.**
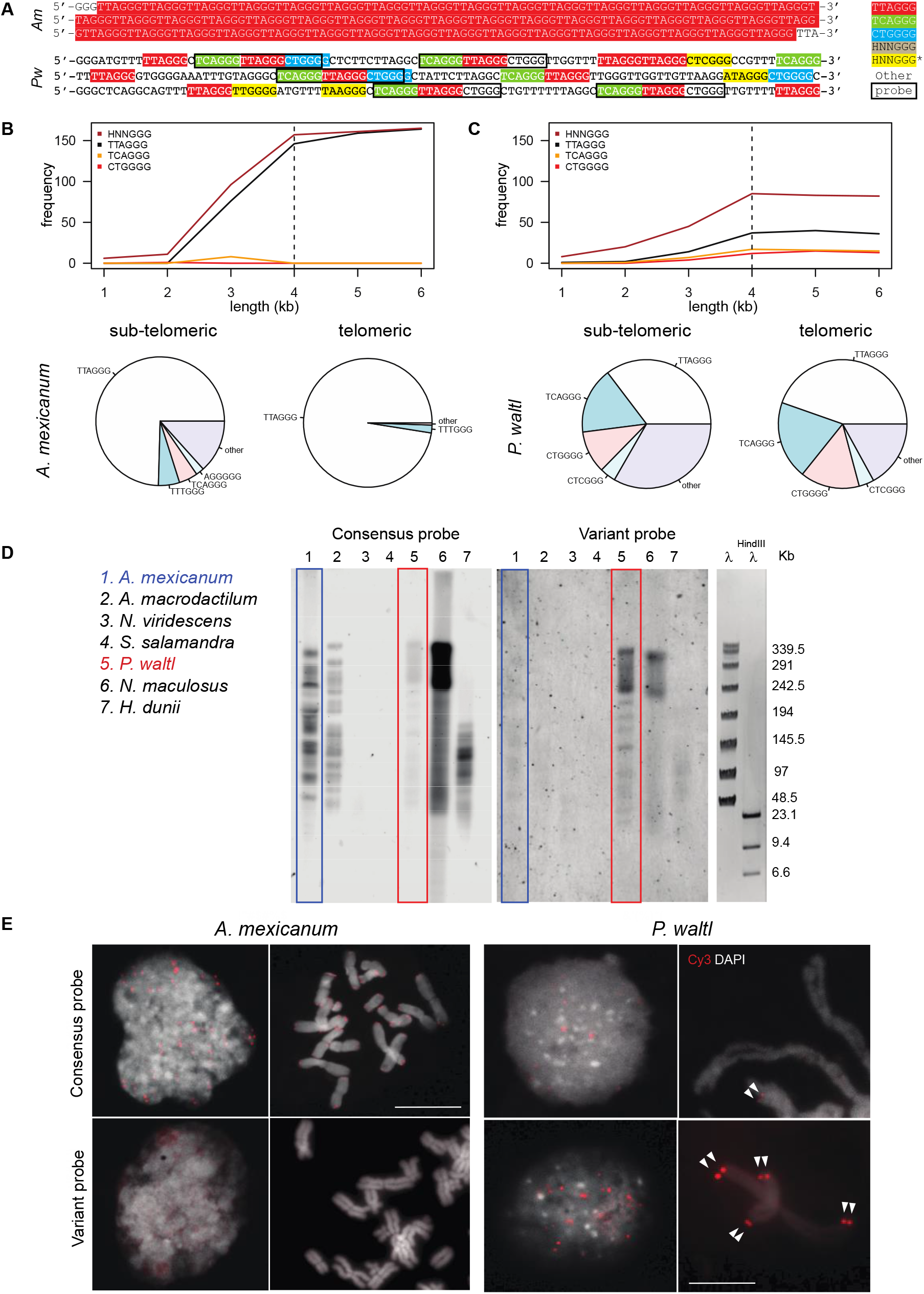
*P. waltl* telomeres consist of interspersed variants repeats. (**A**) Representative 300bp-telomeric region from PacBio genomic data for *A. mexicanum* (top) and three combined Illumina reads for *P. waltl* (bottom). Variant repeats are highlighted in different colours. The ‘variant’ probe used for subsequent analysis is demarcated in black. (**B** and **C**) Top: frequency of telomere repeats in representative telomeric PacBio reads for *A. mexicanum* (B) and *P. waltl* (C). Sub-telomeric and telomeric regions are demarcated by a dotted line. Bottom: Percentage of each telomeric repeat type in the sum of sub-telomeric and telomeric reads for each species. (*) HNNGGG was quantified to exclude G in the first position and prevent counting overlapping hexamers. (**D**) TRF analysis of equal amounts of genomic DNA from the indicated species. The membrane was hybridised with a telomere consensus probe (left), stripped and re-hybridised with the variant probe (right). Note appearance of telomere fragment signal in *P. waltl* upon hybridisation with the variant probe. (**E**) FISH assay using telomere consensus or variant probes on metaphase spreads derived from *A. mexicanum* (left) and *P. waltl* (right) embryonic cells (n=4). Arrowheads highlight telomeric signal in *P. waltl* cells hybridised with the variant probe.

To determine if the *in silico* findings represent genuine chromosome ends, we designed a variant probe against putative *P. waltl* telomeric sequence (Fig. 2A) based on a commonly occurring 18-mer formed by the three most prevalent telomere repeats in *P. waltl* (Fig. S7). Telomere restriction fragment (TRF) analysis of blood samples of seven related species (Fig. 1F) revealed banding patterns consistent with canonical telomeres of the expected molecular weights for members of the *Ambystomatidae, Proteidae*, and *Hynobiidae* salamander families. Minimal to no canonical telomere signal was detected in samples from the newt species (Fig. 2D). Hybridizing with the variant probe, however, revealed a banding pattern consistent with telomeres in *P. waltl* (Fig. 2D) and was further supported by dot blot assays (fig. S3, B and C). Treatment with BAL-31 exonuclease, either alone or in combination with NotI endonuclease (fig. S8, A and B) to expose internal DNA regions to BAL-31 digestion, resulted in a similar reduction of hybridization by the variant probe to isolated *P. waltl* DNA (Fig. S8A). Further, BAL-31 treatment reduced the relative content of *P. waltl* telomeric signal as assessed by qPCR, whereas the interstitial sequence *cdkn2b* remained unaffected (fig. S8C), supporting the terminal nature of the variant repeats. To test this further, we performed FISH on *A. mexicanum* and *P. waltl* metaphase spreads using variant or consensus probes (Fig. 2E and fig. S9). While *A. mexicanum* telomeres were exclusively detected using the consensus sequence, punctate signals at *P. waltl* chromosome termini were readily identified with the variant probe, while no interstitial staining was detected. This provides diret evidence of the terminal nature of the variant repeats. We note the consensus telomere probe reveals weak signals in *P. waltl* FISH and TRF samples (Fig. 2, D and E), consistent with interspersion of approximately 40% canonical hexamers within the abundant variant repeats in this species. Collectively, these data demonstrate that *P. waltl* telomeres lack tandem arrays of consensus repeats and, instead, consist primarily of interspersed variants, a unique feature in vertebrate biology.

Telomere protection is mediated by recruitment of the shelterin complex to chromosome ends(*18*). To determine if salamander telomeres recruit shelterin-components, we expressed GFP-tagged alleles of axolotl or *P. waltl* TRF1 or TRF2 in the corresponding cells. Discrete GFP foci were present in both species and co-localized with telomere sequences as determined by FISH co-staining (Fig. 3, A and B, and fig. S10). Notably, we observed significantly fewer GFP foci in *P. waltl* cells as compared to *A. mexicanum* despite the species containing 12 and 14 chromosome pairs respectively (Fig. 3, A and B). The reason for this discrepancy remains unknown but could reflect telomere clustering in *P. waltl*.

**Fig. 3.**
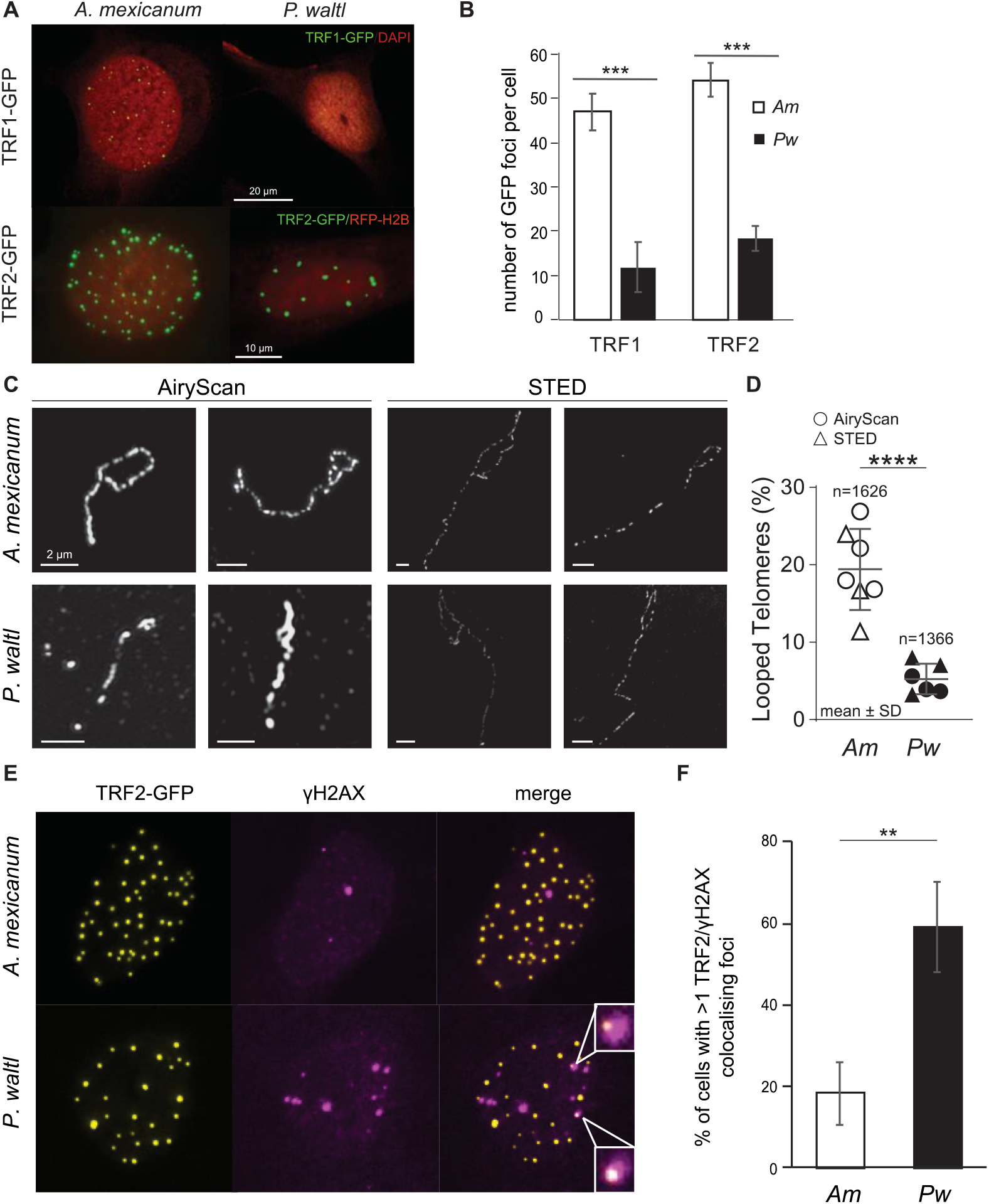
*P. waltl* telomere macromolecular structure is distinct from all known vertebrates. (**A**) Representative images of *A. mexicanum* or *P. waltl* cells 48h after electroporation with pN2-TRF1-GFP or pN2-TRF2-GFP constructs (n=5). **(B)** Quantification of GFP foci per cell. Values represent mean ± SD; ***p<0.001 (n=3; 40 cells scored per replicate). (**C**) Representative images of t-loops from *in situ* trioxsalen crosslinked genomic DNA spread onto glass coverslips and imaged with AirySan or STED super-resolution microscopy. Telomeric DNA was detected by FISH using the consensus or variant probe in *A. mexicanum* and *P. waltl*, respectively. (**D**) Quantitation of the imaging from (**C**) (*A. mexicanum* and *P. waltl* telomeres were quantified from 7 biological replicates each using two different imaging modalities as indicated, **** p < 0.0001. (**E**) Representative images and (**F**) foci quantification of salamander cells 48h post-electroporation with pN2-TRF2-GFP and subsequent immunocytochemistry against γH2AX. Data represent mean ± SEM; **p < 0.01 (n=3; ≥ 20 cells scored per replicate).

Mammalian telomeres are protected by sequestering the chromosome terminus within a lariat conformation called a t-loop(*18*). This structure is conserved across diverse eukaryotic phyla, and its formation in mammalian somatic cells is dependent on TRF2(*19*). We used super-resolution microscopy to investigate macromolecular telomere structure in axolotl and *P. waltl* genomic DNA stained with canonical or variant probes (fig. S11). Canonical axolotl telomeres displayed t-loops consistent with prior observations from other vertebrate species. In contrast to all multicellular organisms surveyed to date(*18*), looped telomeres were present at strikingly low frequency within *P. waltl* telomeres (Fig. 3, C and D, and fig. S11). Sample cross-linking efficiency was equivalent between species (fig S11, B and C) and comparable to that observed in mammals(*20*), and results concordant across super-resolution modalities (Fig. 3D), indicating a bona fide difference between *A. mexicanum* and *P. waltl*. Additionally, we observed an increase in the DNA damage response marker γ-H2AX in *P. waltl* telomeres (Fig. 3, E and F), indicating compromised protection at newt linearised telomeres(*20*).

*P. waltl* telomere composition and structure are reminiscent of that found in human ALT^+^ cells(*15, 16*) and, further, linearized telomeres represent substrates for homology-directed telomere synthesis(*21*). This prompted us to address whether *P. waltl* employs the ALT pathway for telomere maintenance. Indeed, we observed that both FANCD2, a factor recruited to telomere variants which initiates ALT by loading polδ3(*22*), and polδ3 itself, the principal DNA polymerase component underlying ALT(*21*), are actively recruited to telomeric foci in *P. waltl* (Fig. 4, A and B) as previously observed in mammalian ALT contexts(*21*). Similar co-localizations were rare in *A. mexicanum*. Additionally, we observed EdU incorporation in *P. waltl* telomeres (fig. S12) consistent with previous reports of ALT cells displaying elevated telomere synthesis outside of S-phase(*21*). *P. waltl* also contained detectable levels of t-circles, a subset of extrachromosomal telomeric DNA circles that correlate with ALT activity(*23, 24*), in both homeostatic and regenerating tissues (Fig 4, C and D, fig. S13). While the increase in t-circles seen in *P. waltl* is less pronounced than that commonly found in ALT^+^ cancer lines, this may reflect a difference between a physiological context in which the production of deleterious extrachromosomal DNA is controlled (newts), and a deregulated pathological process (cancer cells). Together, these data demonstrates that *P. waltl* exhibits molecular features of ALT activity.

**Fig. 4.**
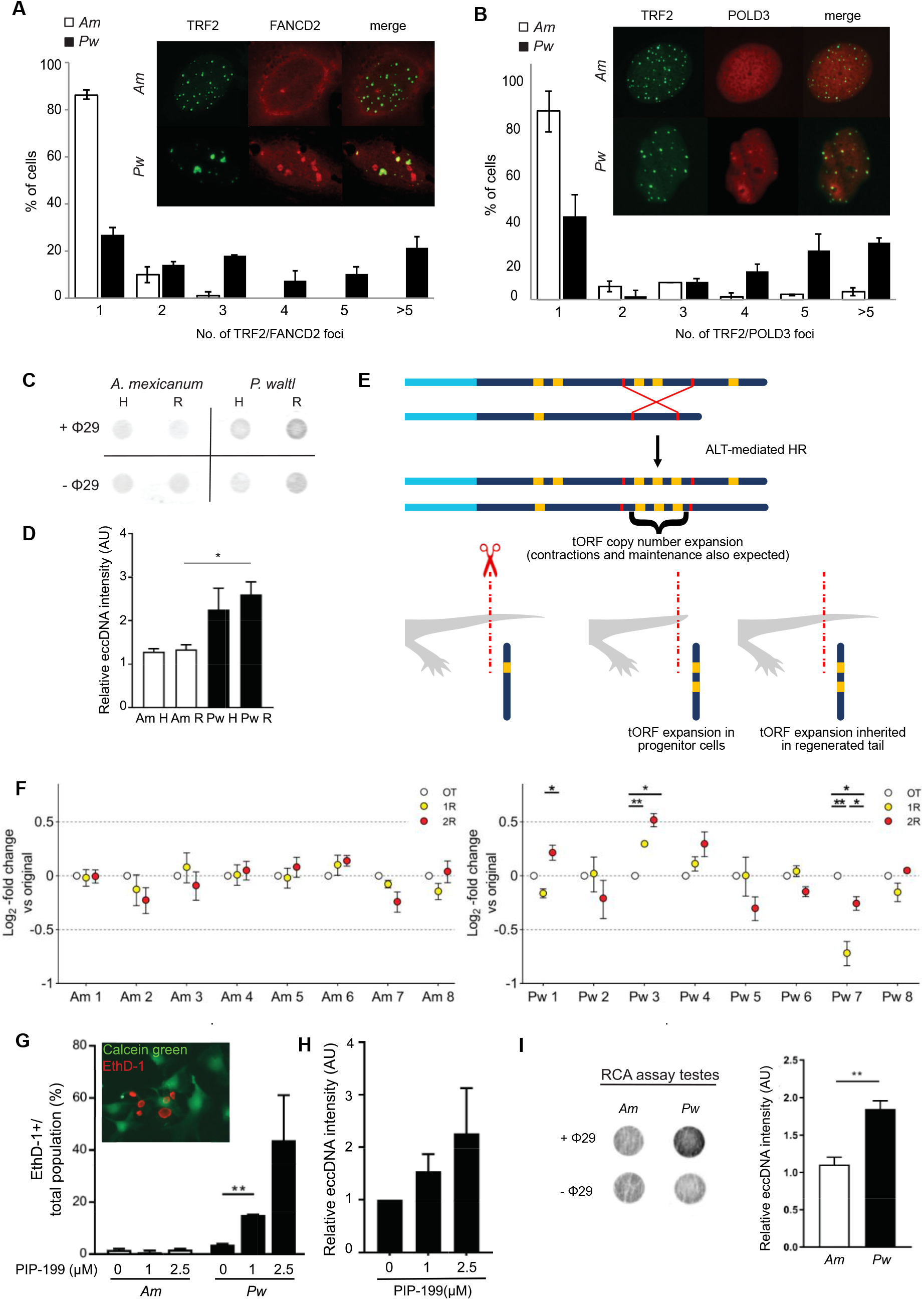
ALT activity in germline and regenerating tissues of *P. waltl*. (**A**) FANCD2 and (**B**) Polδ3 recruitment to *P. waltl* telomeres. Representative images (top) and foci quantification (bottom) of salamander cells 48h post-electroporation with pN2-TRF2-GFP and subsequent immunocytochemistry against FANCD2 (A), or co-electroporation with pN2-polδ3-RFP (B). (n=3; 40 cells scored per replicate). (**C**) Quantitation of t-circle generation during *P. waltl* regeneration. Top: Representative t-circle assay on paired-matched homeostatic (H) and regenerating tail (R) samples for the indicated species (n=3). Dot blots were hybridised with variant (*P. waltl*) or consensus (*A. mexicanum*) probes. -ϕ29 indicates basal DNA signal of the corresponding sample prior to rolling circle amplification. (**D**) Quantification of (C). Data represents eccDNA signal (+ϕ29) relative to background (-ϕ29). (**E**) An approach to estimate ALT activity *in vivo*. Schematic representation of telomeric ORF (tORF) expansion through ALT-mediated recombination (top) and predicted changes through one round of tail regeneration in a given individual (bottom). tORF consists of a 350bp sequence located exclusively within variant repeats. (**F**) Active recombination in *P. waltl* telomeres through serial regeneration rounds. Changes in tORF copy number assessed by qPCR throughout multiple rounds of regeneration in *A. mexicanum* (Am, left) or *P. waltl* (Pw, right) individuals. Results are expressed as fold-change in tORF copy number of first-(1R) and second-regenerated (2R) tails compared with the original (OT) and normalised to genomic *prod1* or *cdkn2b* using the ^ΔΔ^CT method. Data represent mean ± SEM of four reactions with 8 individuals per species. Statistical significance was evaluated by one-way repeated measure ANOVA followed by Tukey’s *post hoc* test. *p<0.05, **p<0.01. (**G** and **H**) PIP-199 induces selective toxicity and increased C-circle generation in *Pw* cells. (G) Quantification of cell death after PIP-199 treatment. (H) Quantification of t-circle assay (n=3). (**I**) T-circle generation in the germline of *P. waltl*. Representative blot (left) and quantification (right) of RCA assay on testes samples for the indicated species (n=6).

Next, we sought to directly assess telomeric recombination events *in vivo* (Fig. 4E). Molecular assessment of ALT-dependent telomere recombination using standard means is hindered in amphibians due to the lack of highly proliferative cell culture. To overcome this hurdle, we developed a novel method to measure telomere recombination *in vivo* through quantitation of the open reading frames (ORFs) present within vertebrate telomeres(*25, 26*). We identified a subset of ORFs present in telomeric regions from *P. waltl* and *A. mexicanum* (designated as tORF). Their terminal nature is further indicated by sensitivity to BAL-31 treatment (fig. S8C). As such, they constitute a genetic tag to evaluate telomeric recombination events. ALT-mediated recombination should result in stochastic tORF copy number variation through expansion, maintenance or contraction (Fig. 4E). Consequently, quantitation of tORFs before and after a replication challenge such as tail regeneration within a single individual offers an estimate of telomeric recombination (Fig. 4E). In contrast, telomere maintenance through telomerase should not lead to tORF copy number alterations. As expected, no significant changes in tORF copy number are observed during tail regeneration in telomerase-dependent axolotls, while extensive changes are observed within successive tails from the same *P. waltl* individuals (Fig. 4F). Further, no significant changes in copy number of a non-telomeric *P. waltl* sequence are observed (fig. S14). These results demonstrate that *P. waltl* telomeres are subject to active recombination. The stochastic nature of this process should result in heterogeneity in telomere size and composition between individuals, as seen in ALT^+^ cells(*24*). In agreement, *P. waltl* tissues displayed TRF heterogeneity, manifest as band smearing, in comparison to the more discrete banding patterns found in axolotl TRFs (fig. S15). This is consistent with the variant repeat heterogeneity exhibited across PacBio telomeric reads in *P. waltl* (fig. S6 and Methods).

Mammalian ALT activity is dependent on FANCM-BTR, and targeting this complex causes selective toxicity in ALT cells through increased telomere replication stress(*27*). We sought to obtain functional confirmation of ALT usage by treating *A. mexicanum* or *P. waltl* cells with PIP-199, a specific FANCM-BTR inhibitor(*27*). *A. mexicanum* cells were not impacted by treatment with PIP-199, while equivalent treatment of *P. waltl* cultures resulted in cell death (Fig. 4G) and a moderate increase in t-circles (Fig. 4H) as previously reported for ALT^+^ cells(*27*).

Critically, ALT is defined as telomere length maintenance in the absence of telomerase. In agreement, we observed telomere length maintenance in three successive *P. waltl* generations (fig. S16) despite no evidence of a *Tert* gene. This is accompanied by the fact that *P. waltl* exists as a species. Within the germline, *P. waltl* testes displayed a greater abundance of t-circles than the equivalent tissue from *A. mexicanum* (Fig. 4I and fig. S17). Further, while axolotl testes displayed conspicuous telomerase activity, no telomerase activity was observed in TRAP assays of *P. waltl* testes using consensus or variant telomere primers (figs. S18 and S19). Collectively, the data are consistent with ALT activity in *P. waltl* reproductive, adult, and regenerative tissues. Therefore, we conclude that *P. waltl* uses the ALT pathway to maintain telomere length throughout lifespan (fig. S20).

Our findings demonstrate that *P. waltl* telomere biology is unique amongst *Chordata. P. waltl* constitutes the first vertebrate species exhibiting telomeres defined primarily by variant repeats, a paucity of t-loops, lack of *Tert* gene, and telomere length maintenance during homeostasis, regeneration and across generations in the absence of telomerase. Although an ALT-dependent strategy has been postulated in a few vertebrate contexts including marsupials(*28*) and bats(*6*), direct proof has been lacking until now. Our findings place *P. waltl* as the first vertebrate species demonstrated to exclusively use ALT at the whole-organism level. How this mechanism evolved and how it is regulated to avoid adverse physiological consequences(*10*) merit further investigation. In this connection, our data indicates that salamanders have long telomeres, suggesting that the ancestor of *Salamandridae* did as well, allowing a larger time window for adaptation to ALT-dependent telomere extension. The prevalence of variant telomere repeats in *P. waltl*, far higher than reported for ALT^+^ cells (> 50% vs <3%)(*16,17*), and is consistent with evolutionary divergence in a species that lost telomerase and gained ALT as the main telomere length maintenance mechanism. Further, the paucity of t-loops in *P. waltl* may indicate a divergence in telomere protection mechanisms which renders telomeres more prone to recombination. Interestingly, failure to detect canonical telomere signal in other *Salamandridae* members suggests these features may be common to the family. As the ability to regenerate complex structures is found across all salamander families regardless of their telomere maintenance mechanism, it is thus unlikely that regeneration is tied to the acquisition of ALT.

Together, our results provide a new model for investigating the mechanisms underlying ALT activity and its regulation, and offer a vertebrate system for *in vivo* screening of ALT inhibitors and compounds of therapeutic potential(*2, 29*). As such, these findings have important implications for our understanding of genome maintenance and its forthcoming applications.

## Acknowledgments

We thank all members of the Yun Lab for critical comments, the Stewart Lab for access to PFGE equipment, Anna Poetsch for experimental advice, Jeremy Brockes for critical reading of the manuscript, Osvaldo Chara for statistical advice, Carlos Talavera-López for processing of raw PacBio subreads, Gabriel Waksman for institutional support, and Beate Gruhl and Anja Wagner for animal care and breeding. The TUD-CMCB Light Microscopy Facility, the CMRI Australian Cancer Research Foundation Telomere Analysis Centre, and the University of Sydney Microscopy & Microanalysis Facility are acknowledged for imaging acquisition and analysis support.

## Author contributions

PBG, QY, SR, IM, AC, ML, NFR and MHY performed experiments. PBG, QY, SR, IM, ML, FS, AE, AC, AS and MHY analysed and discussed data. PBG, QY, IM and MHY designed experiments. MHY supervised the project and wrote the manuscript with contributions from all authors.

## Data and materials availability

*P. waltl* PacBio subreads (fastq and fasta files) are available under BioProject PRJNA353981 (SRR8594170). All other data is available in the main text or the supplementary materials.

## Supplementary Materials

Materials and Methods

Supplementary Text

Figs. S1 to S19

Tables S1 to S3

References (*31*–*39*)

