## supplemental information for "Telomerase-independent maintenance of telomere length in a vertebrate"

**This PDF file includes:**

Materials and Methods
Supplementary Text
Figs. S1 to S20
Tables S1 to S3

### Materials and Methods

Animal husbandry

Procedures for care and manipulation of all animals used in this study were performed in compliance with the Animals -Scientific Procedures- Act 1986 (United Kingdom Home Office), and the laws and regulations of the State of Saxony, Germany. *Pleurodeles waltl* (*P. waltl*) newts were obtained from the Newt Facility at TUD-CRTD Center for Regenerative Therapies Dresden (Germany). *N. viridescens* newts were obtained from Charles Sullivan and Co. (Tennessee, USA) and maintained as described elsewhere (*31*). Axolotls (*A. mexicanum*) were obtained from Neil Hardy Aquatica (Croydon, UK) and from the axolotl Facility at TUD-CRTD Center for Regenerative Therapies Dresden (Germany). *A. macrodactilum*, *S. salamandra* and *H. dunii* specimens were obtained from Neil Hardy Aquatica (Croydon, UK). *N. maculosus* specimens were obtained from the biological facility at University College London (UK). Axolotls were maintained in individual aquaria at approximately 18°C, as previously described (*10*). *P. waltl* were maintained in group holding at 20°C, as described (*32*). All other salamander specimens were processed for sample collection immediately upon acquisition.

Animal procedures and tissue collection

Newts and axolotls were anesthetised in 0.1 % tricaine prior to limb amputation at the mid-humerus level, or amputation of the last third of the tail. Animals were allowed to regenerate at 20°C and tails, limbs or blastemas collected and processed for various assays as specified. Axolotl regeneration stages were defined as previously described (*33*).

For telomere restriction fragment analysis (TRF), animals of comparable age (2-3 years old) were anaesthetised and terminated. Immediately, blood samples were collected into 5 ml 10 mM EDTA/A-PBS (PBS + 25 % $dH_2O$), spun down at 1000 rpm for 5 minutes, washed in A-PBS, and embedded in agarose plugs as described below. For TRAPeze and C-circle assays during spermatogenesis, testes were collected from one-year old sexually-mature *A. mexicanum* and *P. waltl* males, washed with 0.8 X PBS, and fast-frozen in liquid nitrogen. One testis from each animal was used for TRAPeze assays and the contralateral testis processed for C-circle assays. Freshly-hatched 3-week-old larvae were collected as above.

*A. mexicanum* developmental stages were determined according to (*34*). *P. waltl* developmental stages were determined as specified in (*32*).

Cell culture

A1 cells were previously derived from newt limb mesenchyme (*31*). Axolotl AL1 cells were obtained from S. Roy (University of Montreal, Montreal, QC, Canada). *P. waltl* primary cells derived from embryonic tissues were obtained from P. Ferretti (University College London, UK). Pwl1 cells were derived from newt limb connective tissue, grown for three months as described below and subsequently expanded. A1, AL1, Pwl1 and *P. waltl* primary cells were grown on 0.75% gelatin-coated plastic dishes in MEM (Gibco, UK) supplemented with 10 % heat-inactivated foetal calf serum (FCS, Gibco), 25 % H2O, 2 nM L-Glutamine (Gibco), 10 µg/ml insulin (Sigma, St Louis, MO) and 100 U/ml penicillin/streptomycin (Gibco) in a humidified atmosphere of 2.5 % CO2 at 25°C. Cell subculture was performed as previously (*35*). For the generation of primary spleen cultures, whole spleens were isolated, mechanically dissociated by pipetting in 1X trypsin for 20 minutes and cultured in L15 (GIBCO) supplemented with 10 % heat-inactivated foetal calf serum (FCS, Gibco), 25 % H2O, 2 nM L-Glutamine (Gibco) and 100 U/ml penicillin/streptomycin (Gibco), in addition to 100 µg/ml colcemid for 2 days.

Telomerase activity assay

Telomerase enzymatic activity was assessed using the TRAPeze telomerase assay kit (MERCK), according to manufacturer's instructions. Samples were homogenized in 1X CHAPS lysis buffer. Tissue homogenates were incubated for 30 minutes on ice and then centrifuged for 20 minutes (12,000 *g*, 4°C). Protein concentrations of homogenates were quantified using the BCA method and 50 ng of total protein was used for each reaction. Indicated samples were incubated at 85°C for 10 minutes for heat inactivation. Telomerase extension was performed at 30°C for 30 minutes, followed by PCR amplification of extension products. End products were resolved by PAGE [pre-cast TBE 10% gels (Thermo), 200 V, 50 minutes in XCell SureLock Mini-Cells (Thermo)], stained with SYBR Safe Nucleic Acid gel stain for 30 minutes at room temperature, and visualized on a Typhoon 9210 scanner. TRAPeze assays with alternative reverse primer (variant-RP) experiments were canonical TRAPeze assays, but with the canonical RP substituted for a primer designed for extension product amplification based on the most frequent *P. waltl* telomeric motif (5'-CCCTGACCCAGCCCTAACCCTGA-3').

*Tert* gene/transcript mining

Protein sequences for *X. laevis* and *X. tropicalis Tert*, *A. mexicanum Tert* and the genes surrounding *Tert* in the axolotl genome assembly (*11*) *Top*1.2, *Clch*1, *Rps*27a, *Mtif*2.L, *Mlycd* (Table S1) were queried using tblastn against the *P. waltl* and *N. viridiscens* transcriptomes (*12*) with an evalue cutoff of 0.1 (Table S2). Xenopus and axolotl TERT protein sequences were also queried using tblastn against the *P. waltl* genome assembly (*12*) with an evalue cutoff of 0.1. The coordinates for the TERT locus in the axolotl genome are AMEXG_0030001724:5443375-6406992 (assembly ambMex 3.0.0), accessible at https://genome.axolotl-omics.org/.

Identification of telomeric sequences in *A. mexicanum*

The axolotl genome sequence ((*11*)PMID: 29364872) in FASTA format was queried using the grep command to retrieve contigs with hexamers TTAGGG or CCCTAA repeated at least 5 times. This step resulted in 39 contigs. Sequences were manually inspected to exclude contigs with blocks of TTAGGG repeats present in non-terminal regions (interstitial telomeric sequences) and contigs with blocks of TTAGGG repeats throughout the entire span (i.e. without sub-telomeric regions). The remaining five contigs were selected to represent axolotl telomeric and subtelomeric sequences for Fig. S6A.

Identification of telomeric sequences in *P. waltl*

*P. waltl* telomeric sequences were retrieved from a published WGS Illumina library (SRR6001099, (*12*)) and PacBio subreads of the *P. waltl* genome (see 'Resource sharing' section). The genomic DNA sequenced using PacBio originated from the same sample described in (*12*). PacBio library preparation and sequencing was done at SciLifeLabs (Uppsala). Individual tracks of hexamers CTAGGG, CTTGGG, TCAGGG, TTAGGG, TTCGGG, TTGGGG or TTTGGG, each repeated 110 times were aligned using blastn to PacBio subreads (~ 72Gb) of the *P. waltl* genome. To retrieve telomere sequences, blastn was used without filtering low complexity regions in the query and database (-dust no -soft_masking false). The penalties for opening and extending an alignment gap or for a mismatch were adjusted to accommodate the erroneous indels and mismatches found in uncorrected PacBio subreads (-gapopen 0 -gapextend 2 -penalty -1). Finally, the length of the initial exact match was reduced to 6 bases, in line with the telomere repeat being a hexamer (-word_size 6).

PacBio subreads aligning to more than four of the query tracks (349 reads) were manually inspected to exclude subreads with blocks of telomeric repeats present in non-terminal regions (interstitial telomeric sequences) and subreads with blocks of telomeric repeats throughout the entire span (i.e. without sub-telomeric regions). The remaining ten subreads were selected to represent newt telomeric and subtelomeric sequences for Supplementary Figures S6(B).
The repetitive element TCAGGGTTAGGGCTGGGG was identified as a recurrent variant telomeric repeat using PacBio reads (see below). To confirm that the variant repeats are not an artifact of PacBio sequencing errors, the element TCAGGGTTAGGGCTGGGGTCAGGG TTAGGGCTGGGG (i.e. TCAGGGTTAGGGCTGGGG repeated twice) was aligned to the Illumina library SRR6001099 using the online NCBI BLAST interface and retrieved 100 hits. Conversely, aligning the consensus telomere repeat TTAGGGTTAGGGTTAGGGTTAGGG TTAGGGTTAGGG to the same library retrieved zero hits. Reads SRR6001099.138434328.1, SRR6001099.152158295.1 and SRR6001099.209958579.1 were used for figure 3A.

Axolotl *Tert* expression
RNA was isolated from axolotl tissues using Tri Reagent (Sigma) and random primed cDNA synthesized using Superscript II (Invitrogen). Gene expression was determined by quantitative real time PCR using the indicated primers (Table S3). RT-PCR was carried out using iQ SYBR Green supermix (BioRad) on a Chromo 4 instrument running Opticon 3 software (Bio-rad). All reactions were run in triplicate and at least 3 independent RNA preparations were analyzed for each sample.

Telomere size calculation
200 mg of salamander forelimb tissue were ground to a fine powder with liquid nitrogen in a percussion mortar and genomic DNA isolated using the blood and cell culture DNA midi kit (QIAGEN) as per manufacturer's instructions. Absolute telomere size for each of the indicated species was determined by quantitative real time PCR on genomic DNA according to (*36*), using the indicated primers and standard oligomers (Table S3). Telomere size was calculated assuming the following diploid numbers: *A. mexicanum* 28, *A. macrodactylum* 26, *P. waltl* 24, *N. viridescens* 22, *S. salamandra* 24, *N. maculosus* 38, *H. dunii* 56.

Metaphase spreads
For cell cultures, cells were trypsinised, washed in A-PBS, resuspended in 0.075 M sodium citrate for 45 minutes at 37°C and fixed by mixing with ice-cold 3:1 methanol:acetic acid drop-by-drop. The resulting cell suspension was then dropped from a distance of 80 cm onto Superfrost slides (20 - 30 µl per slide) and air-dried overnight. For axolotl and newt embryos, eggs were collected at stage 20, dejellied and incubated in 0.1 X MMR (Marks Modified Ringers solution) supplemented with 0.4 % (newt eggs) or 0.25 % (axolotl eggs) colcemid (Sigma) for 18 hours at 20°C. Embryos were fixed in ice-cold 1:1 acetic acid:ethanol for 5 minutes, washed in $H_2O$, transferred to ice-cold 1:1 acetic acid:ethanol and mechanically dissociated by pipetting before dropping the solution onto Superfrost slides as indicated.

Fluorescence in situ hybridization (FISH)
In situ hybridization was carried out in metaphase spreads, containing metaphase chromosomes and interphase nuclei, using the Telomere PNA FISH Kit/Cy3 (Dako) as per manufacturer's instructions. For detection of *P. waltl* telomeres and *N. viridescens* common repetitive element, RE (consisting of consecutive repeats of the sequence 5'-

GGTATAGGTATAGGTATAGGTATA-3’), the corresponding PNA probes (variant probe: Cy3-CCCAGCCCTAACCCTGA-3’, RE probe: Cy5-CCTATACCTATACCTATACC-3’, Biomers) were used in combination with the Telomere PNA FISH Kit/Cy3, in place of the telomere consensus PNA probe (Cy3-OO-TTAGGGTTAGGGTTAGGG-3’, Dako).

EdU/FISH detection and colocalization analysis

Cells were pulsed with 10 µM EdU (5-ethynyl-2’-deoxyuridine) for 2 h, fixed in 3.7% formaldehyde/PBS solution for 10 minutes and permeabilised in 0.2 % Triton X-100/PBS for 15 minutes. The Click-iT Alexa Fluor 488 azide reaction was then carried out according to the manufacturer’s instructions (Invitrogen), before performing telomere staining by FISH as described above. Imaging was performed using a Zeiss LSM 780 confocal microscope, with single plane sequential acquisition of three channels using the same dichromatic mirror (DAPI: ex. 405nm, em. 410-487nm; EdU-Alexa488: ex. 488nm, em. 491-562nm; telomeres-Cy3: ex. 561nm, em. 571-642nm) and a 40x/1,2 W C-Apochromat objective. Images were analyzed using Arivis Vision4D (V 3.1.3). Images were denoised, background corrected and automatically segmented by the blob finder module (telomeres) and the Otsu automated threshold (EdU) module. Complete overlap of telomeres with EdU was calculated using the compartmentalization module.

Telomere restriction fragment analysis (TRF)

DNA was obtained by embedding salamander blood samples from adult animals of comparable age into agarose plugs, followed by an extensive Proteinase K treatment, as described in the CHEF Genomic DNA Plug Kit (Bio-Rad) instruction manual. The plugs were exchanged from 1 ml of 0.1X CHEF wash buffer into 1 ml of 1X CutSmart buffer and incubated for 1 h at RT. The buffer was exchanged for 300 µl of fresh solution and the DNA was digested with 50 units of HinfI, RsaI, HphI and MnlI (New England Biolabs) at 37°C overnight. The plugs were washed with 1 ml of 1X CHEF wash buffer for 1 h at room temperature, cut to 1/10 of the plug size, weighed to ensure uniformity and equilibrated in 1 ml of 0.5 X TBE buffer for 30 minutes at RT. The CHEF DNA size standard (Bio-Rad) was prepared according to manufacturer’s instructions. The plugs were inserted into the wells of a 14x21 cm 1 % agarose/0.5 X TBE gel, sealed with melted agarose and run at 14°C using the CHEF Mapper XA Pulse Field Electrophoresis System (Bio-Rad). The run settings were specified using the auto algorithm option with size range set to 10-400 kb, ramping factor of 0.243 and buffer flow rate of 70. Total run time was automatically determined to be 20:21 h. After the run, the gel was briefly washed with $dH_2O$ and incubated in aqueous solution with 5 µL of SYBR Safe (Invitrogen) for 30 minutes at RT with gentle shaking and de-stained for 5 minutes in $dH_2O$ under the same conditions. The gel was imaged using the Gel Doc EZ Imager (Bio-Rad). It was rinsed in $diH_2O$ and depurinated twice for 15 minutes in a 0.25 M HCl solution, followed by denaturation for 2x20 minutes in 0.5 M NaOH 1.5 M NaCl and neutralization for 2x20 minutes in 1 M Tris 1.5 M NaCl pH 7.5. The rest of Southern blotting was performed as described in Thermo Fisher Scientific instructions, using a Whatman Nytran SPC 0.45 nylon transfer membrane (Sigma). After UV-crosslinking using Stratalinker 1800 at 254 nm (Stratagene) at 160 000 microjoules/$cm^2$, the membrane was prehybridized in 8 ml of 5X SSPE 1 % SDS 0.01 % tetrasodium pyrophosphate 3X Denhardt’s solution for 30 minutes at 37°C. The membrane was hybridized in 7 ml of fresh solution and 1 µL of 100 µM DNA probe (variant probe: 5’-[D3-PA]BYTYAGGGGTTAGGGCT, consensus probe: 5’-[D3-PA]TTAGGGTTAGGGTTAGGGTTAGGG; Sigma Aldrich) at 37°C overnight. It was then washed for 20 minutes at 37°C in 3 X SSC 0.5 % SDS solution and imaged using the Odyssey IR

Imaging System at 700 nm (Li-Cor). For re-probing, membranes were stripped after first detection with 0.2 X SSC 0.5% SDS for 15 minutes at 65°C.

BAL-31 dot blot and qPCR analysis

*P. waltl* DNA was isolated from metamorphosed *P. waltl* limbs using the Genomic-tip 500/G (QIAGEN) and isopropanol precipitation, as described in the QIAGEN Genomic-tip procedure. 200 ng of DNA were incubated with 2 units of BAL-31 exonuclease (New England Biolabs) and 1X BAL31 buffer in a total volume of 20 µl for 1, 2.5, 15, 30 or 60 minutes at 30°C. A subset of samples was pre-digested with NotI-HF (New England Biolabs) for 2 h at 37°C followed by enzyme inactivation for 15 minutes at 70°C. BAL-31 was inactivated with 1 µl of 1 M NaOH 200 mM EDTA for 15 minutes at 75°C. The DNA was then precipitated overnight at -20°C with 70 µl of absolute ethanol, 7 µl of 10 M ammonium acetate and 1 µl of 20 mg/ml glycogen and centrifuged for 30 minutes at 19 000 g, 4°C. Subsequently, 300 µl of cold 70% ethanol were added, and the pellet was re-centrifuged. DNA was air-dried for 20 minutes before dissolving in 20 µl of 10 mM Tris pH 7.6 for 20 minutes at 45°C. Samples were then analysed either through dot blot or qPCR. The dot blotting procedure was performed using a dot-blotter (Carl Roth), as described in (*37*). Hybridization, washing, and detection were performed as described for TRF analysis. Measurement of telomere, tORF, and *cdkn2b* content by qPCR was performed as described for the telomeric ORF recombination assay, with 0.5 ng BAL-31-treated gDNA loaded per reaction and 500 nM primers as indicated in Table S3. Relative content following BAL-31-treatment was calculated using the $2^{-\Delta\Delta CT}$ calculation method, using genomic *e1fα* as reference and normalised against untreated (-BAL-31) samples.

TERT Southern dot-blot analysis

A 500 bp amplicon corresponding to the conserved TRBD domain of TERT was amplified from *A. mexicanum* genomic DNA (forward *tert* primer: CCGAAACGCTATTGGCCCATGAAATG; reverse *tert* primer: CGATACTTGTTATGGTTCGATCC ) and used as a template for digoxigenin-labelled probe synthesis using the random labelling method (DIG High Prime Labelling and Detection Kit, Roche). 5.6 µg *A. mexicanum* and 3.5 µg *P. waltl* gDNA (equivalent to 0.1 pg of target sequence) was blotted onto a Nytran SPC membrane, crosslinked, and hybridized at low stringency (35°C) with 25 ng/ml TERT-digoxigenin probe. Subsequently, membranes were washed at low strignency twice with 2X SSC, 0.1% SDS at 25°C and hybridized probe was visualized using immunological detection followed by colorimetric development with NBT/BCIP for 4 h according to the manufacturer's instructions. Chromogenic product and bound probe were stripped with DMF at 65°C followed by treatment with 0.2M NaOH, 0.1% SDS at 37°C, and then re-probed using D3-labelled consensus and variant telomeric probes as described above to control for gDNA loading.

TRF1 cloning

*A. mexicanum* *Terf*1 was amplified from axolotl normal limb cDNA using *Terf*1 primers: AACCTCGAGCCACCATGGAGGACGTGAACGTGCCATT forward and CGGGAATTCAATGAGATGTAGTTTCTTCATAGTCCTCC reverse.

*P. waltl* *Terf*1 was amplified from normal limb cDNA using *Terf*1 primers: CTCCTCGAGCCACCATGGCCGCCGTTAAAGAGTC forward and GCGGAATTCGCCAGGTATGCTTTGTCTTTGCA reverse. Amplified *Terf*1 sequences were cloned into the XhoI and EcoRI sites of pEGFP-N2 (Clontech).

TRF2 cloning

*P. waltl Terf2* was amplified from 100 ng *P. waltl* spleen cDNA using the primer pair ggtggtGAGCTCATGGCGGGGAGTCAGG forward and tcgtccGAATTCCACCAGTCCTAATCTCTTCATAGTC reverse. The 1.4 kb product was gel purified and subcloned into the pN2-CMV:EGFP transfection plasmid through restriction enzyme digestion with SacI-HF and EcoRI-HF, to generate a PwTRF2-EGFP fusion protein.

Polδ3 cloning

*A. mexicanum Polδ3* was amplified from axolotl larvae cDNA using the primer pair CTCGAGGCCACCATGGACGAGCTTTATCTTGAGAACATAG forward and GGGCGAATTCGGGTTTCTTCTGGAAGAAGCCCATGATG reverse.

*P. waltl Polδ3* was amplified from *P. waltl* larvae cDNA using the primer pair CTCGAGGCCACCATGGACGAGCTGTATCTGGAAAACATCG forward and GGGCGAATTCGGGTTTCTTCTGAAAGAAGCCCATGATGGAG reverse.

The products were gel purified and subcloned into the pN2-CMV:EGFP transfection plasmid through restriction enzyme digestion with XhoI and EcoRI, to generate Pol*δ*3-EGFP fusion proteins.

Cell electroporation

Cells were transfected by nucleofection using the Lonza nucleofection apparatus and reagents. Cell suspensions ($10^5$ cells in 0.1 ml) were mixed with 2 μg pN2-TRF1-GFP or pN2-TRF2-GFP plus 1 μg pN2-RFP-H2B, and nucleofected as per manufacturer's instructions. The nucleofected cells were added to 1.5 ml supplemented minimal medium (MEM, Gibco) and incubated at 25 °C for 20 minutes prior to plating into gelatin-coated dishes. Imaging was carried out at 48 hours post nucleofection.

Immunofluorescence

For staining of cultured cells, these were fixed in 4 % PFA for 5 minutes and processed as described elsewhere(*35*). Fixed cell samples were incubated overnight with anti-γH2AX (S139) mouse monoclonal antibody (Upstate, 1:1000) or anti-FANCD2 rabbit polyclonal antibody (Novus Biologicals, 1:250), and secondary staining was performed using anti-mouse or anti-rabbit AlexaFluor568 antibodies (Invitrogen; 1:1000). Hoechst 33258 (2 μg/ml) was used for nuclei counterstaining. Samples were observed under a Zeiss Axioskop2 microscope and images were acquired with a Hamamatsu Orca camera using Openlab (Improvision) software. Whenever comparative analyses between samples were performed, all images were acquired with identical camera settings and illumination control. Image processing (contrast enhancement) was equally applied to all matched experimental and control samples using Openlab software.

Rolling circle amplification assays

Rolling circle amplification (RCA) assays were performed as previously described(*37*). Briefly, DNA was purified from freshly collected testes, blastema, or homeostatic tissue using the Qiagen DNA Kit according to the manufacturer's instructions. 10 ng of DNA was combined with 0.2 mg $ml^{-1}$ BSA, 0.1 % Tween, 1 mM each dNTP, 1 X Φ29 Buffer (NEB) and 7.5 U Φ29 DNA polymerase (New England Biolabs). Rolling circle amplification was performed for 8 h at 30°C followed by 20 minutes at 65°C. Rolling circle amplification products were then diluted in 2 X

SSC buffer and dot-blotted onto a 2X SSC-soaked Nytran SPC membrane. Membrane was crosslinked and hybridized with 5'-D3-labelled consensus or variant probes overnight at 37°C. Subsequently, membranes were washed for 30 minutes at 37°C and detected using an Odyssey Infrared Imager (Li-Cor).

Telomeric ORF recombination assay

Animals (5-6 cm STL) were amputated at the tail for three rounds of repetitive amputation and regeneration. Tissue from the original, first- and second-regenerated tails were fast-frozen in liquid nitrogen and stored at -80°C until processing. Genomic DNA was extracted with DNeasy Blood and Tissue Kit (Qiagen) according to the manufacturer's instructions and quantified in triplicate by a NanoDrop 2000 spectrophotometer (Thermo). 0.5 ng gDNA was loaded per reaction with 1X SybrGreen Supermix (BioRad) and 500 nM forward and reverse primers as indicated (Table S3). Relative copy number of tORF was measured using a CFX Connect RT-PCR machine (BioRad). For each species, tORF and *cdkn2b* (*P. waltl*) or *prod1* (*A. mexicanum*) qPCRs were performed in parallel, on the same plate, in duplicates. Each assay was performed in four replicates. The relative tORF copy number was normalized to *cdkn2b* or *prod1* according to the $2^{-\Delta\Delta CT}$ calculation method. qPCR was performed in four replicates with two technical repeats per sample in each run. Statistical analysis was performed with one-way repeated measures ANOVA followed by Tukey's post-hoc test.

Toxicity assay

AL1 or Pwl1 cells were plated at 10,000 cells/24-well plate and 24 h afterwards treated with 0, 1 or 2.5µM PIP-199 in dimethyl sulfoxide (DMSO). Drug or vehicle were administered daily for a period of 6 days. Cell death and survival were estimated at end point by incubating cells in 1.5µM Ethidium Homodimer-1 (dead cell stain, Invitrogen) and 150nM Calcein Green AM (live cell stain, Invitrogen), followed by live imaging in an AxioZoom V16 fluorescent stereoscope (Zeiss). For each replicate, parallel cultures were set up for RCA assays, performed as described above.

Isolation and crosslinking of nuclei extracted from frozen tissue for t-loop assays

The following steps were adapted from Corces et al.(*38*). Briefly, 50 mg of A. mexicanum and P. waltl liver tissue were cut from snap-frozen sections and allowed to thaw in 2 mL ice-cold homogenization buffer [20 mM Tricine-KOH pH 7.8, 260 mM Sucrose, 30 mM KCl, 10 mM $MgCl_2$, 500 µM Spermidine, 150 µM Spermine, 1 mM DTT, 0.3% NP40, cOmplete Protease Inhibitor (Roche)], before homogenization in a prechilled glass dounce for 10x strokes with a loose pestle, and 20 strokes with a tight pestle. Lysates were filtered using a 70 µm cell strainer and centrifuged at 350 rcf for 10 min in LoBind 5 mL tubes to pellet nuclei. The extracted nuclei were resuspended to a volume of 400 µL in homogenization buffer and then mixed with an equal volume of 50% iodixanol solution [60% iodixanol solution (Sigma) diluted with 120 mM Tricine-KOH pH 7.8, 150 mM KCl, 30 mM $MgCl_2$]. Layers of 30% and 40% iodixanol lysis solution (made from 50% iodixanol solution diluted in lysis buffer) were layered underneath the nuclei containing fraction. The layered fractions were centrifuged at 600 rcf for 20 min without brakes. Purified nuclei were pipetted from the 30%-40% iodixanol interface, diluted with nuclei wash buffer (10 mM Tris-HCl pH 7.4, 15 mM NaCl, 60 mM KCl, 5 mM EDTA, 300 mM Sucrose), and pelleted at 350 rcf for 10 min. Approximately $5e^5$ nuclei were resuspended in 1 mL nuclei wash buffer containing 100 µg/mL Trioxsalen (Sigma) (diluted to 5 mg/mL in DMF). The remaining steps were adapted from Van Ly et al.(*20*). The nuclei solution was exposed to 365 nm UV light 2-3 cm from the irradiator (model UVL-56, UVP) for 30 min inside a non-TC treated 6 well plate cooled on ice with constant stirring. Crosslinked nuclei were centrifuged at 350 rcf for 10 min and washed with nuclei wash buffer. Material was resuspended in nuclei wash buffer and mixed 1:10 with spreading buffer (10 mM Tris-HCl pH 7.4, 10 mM EDTA, 0.05% SDS, 1 M NaCl) prewarmed to 40 °C. 100 uL of the chromatin solution was immediately spun onto 18x18 mm, 0.17 mm thick acid-washed glass coverslips using a Cellspin1 (Tharmac) at 600 rpm for 1 min. DNA-coated coverslips were fixed in -20 °C anhydrous methanol for 10 min, followed by -20 °C acetone for 1 min. The fixed coverslips were briefly rinsed with PBS, followed by deionised water, and then dehydrated though a graded series of ethanol (70, 90, 100%) for 5 min each.

Telomere fluorescent *in situ* hybridisation (FISH) for AiryScan and STED

Ethanol dehydrated coverslips were denatured for 10 min at 80 °C in the presence of C-rich telomere peptide nucleic acid (PNA) probe conjugated to Alexa Fluor 647 (Alexa647-OO-CCCTAACCCTAACCCTAA, Panagene) for *A. mexicanum* AiryScan and STED imaging, or Cy3 (Cy3-OO-CCCAGCCCTAACCCTGA, Panagene) for *P. waltl* AiryScan imaging, and Alexa Fluor 568 (Alexa568-OO-CCCAGCCCTAACCCTGA, Panagene) for *P. waltl* STED imaging. PNA probe was prepared and diluted to 0.3 ng/mL as described previously(*39*). Following hybridization overnight in a dark humidified box, coverslips were washed twice for 10 min in PNA Wash A (70% Formamide; 10 mM Tris-HCL pH 7.5) and thrice for 5 min in PNA Wash B (50 mM Tris-HCl pH 7.5, 150 mM NaCl, 0.8% Tween-20), all with gentle shaking, and counterstained with DAPI (AiryScan) or YoYo-1 (STED). Coverslips were rinsed in MilliQ water and dehydrated through a 70%, 90%, and 100% ethanol series. Coverslips were air-dried, stored dehydrated, and then mounted using Prolong Gold (AiryScan, Life Technologies) or Diamond (STED, Life Technologies) before imaging.

AiryScan and STED Imaging

AiryScan imaging was performed on a ZEISS LSM880 AxioObserver confocal fluorescent microscope fitted with an AiryScan detector and a Plan-Apochromat 63× 1.4 NA M27 oil objective. Chromatin was identified with DAPI using conventional resolution microscopy prior to AiryScan imaging. Alexa Fluor 647 labeled telomeres were captured using the 633 nm laser at 18% transmission power, 1×1 binning, detector gain of 906 and digital gain of 1.0 in super-resolution mode. Cy3 labeled telomeres were captured using the 561 nm laser at 0.2% transmission power, 1×1 binning, detector gain of 906 and digital gain of 1.2 in super-resolution mode. A total of 5 z stacks (200 nm) were captured with frame scanning mode, unidirectional scanning, and line averaging of 2 in 1024 × 1024 pixels at 36.61 x 36.61 μm to scale. Z stacks were AiryScan processed in Zen Black software (ZEISS). Images were scored, measured and adjusted using FIJI/ImageJ (1.53c), and arranged using Adobe Illustrator (24.3.2) and shown here, as cropped maximum intensity projections.

STED imaging was performed on a Leica TCS SP8 microscope (Leica Microsystems GmbH, Mannheim, Germany), using a 93x 1.30 NA glycerol immersion motCORR STED White objective and a tunable white light laser unit. Alexa Fluor 647 labeled telomeres were imaged with a 647 nm excitation running at ~12% power, and with emission wavelengths set between 660 and 750 nm. Fluorescence depletion used a 775 nm laser running at 22.5% output power. Pinhole was set at 152 μm. For telomeres stained with Alexa Fluor 568, imaging was performed using a 568 nm excitation line set at 15-20% power and an emission wavelength range of 588 nm - 702 nm. The 775nm STED depletion laser was set 95%. A pinhole size of 92 μm was used. Both emitted fluorescence intensities were filtered by a notch filter (775 nm). Images (1024 by 1024 pixels) were collected with a pixel size range of ~ 25 nm – 50 nm (Zoom range between 2 to 6) and with a Z-stack step size of 0.1 μm (typical range of 0.5 – 1.0 μm). A line average of 4 and a frame accumulation of 2 was used for the 647 nm channel; a line average of 6 with a frame accumulation of 2 or 3 was used for the 568 nm labeled samples. All scans were performed at a scan speed of 600 Hz. Deconvolution of images was performed with Huygens Professional software (Scientific Volume Imaging, Hilversum, The Netherlands). Images were scored and measured using FIJI/ImageJ (1.53c), levels were minimally adjusted in Adobe Photoshop (22.3.1) and shown here, as cropped maximum intensity projections.

We estimated microscope resolution using an intensity profile generated by PlotProfile (ImageJ) from a line scan across DNA fibres within maximum projection images. Full width at half maximum height (FWHM) was calculated using σ derived from a gaussian fit curve, and the equation:

$$FWHM = 2\sqrt{2\ln(2)} * \sigma$$

These calculations produced the following resolutions (mean ± SD, n=9): AiryScan Cy3 : 171 ± 20 nm; AiryScan Alexa647 : 177 ± 12 nm ; STED Alexa568 : 112 ± 37 nm; STED Alexa647 : 91.5 ± 36 nm.

Crosslinking Efficiency Determination

The nuclei extraction protocol for super-resolution microscopy was followed through the trioxsalen crosslinking step on ~600 mg of tissue. Crosslinked nuclei and matched non-crosslinked control nuclei were pelleted by centrifugation at 350 g for 10 min at 4 °C. The DNA was digested with 60 U/1e$^{6}$ nuclei of micrococcal nuclease (NEB) at 37°C for 10 min. The reaction stopped with equivolume ice-cold nuclei wash buffer and the chromatin pelleted at 1500 g for 5 min. Chromatin pellets were resuspended in 70 mM Tris-HCl pH 8.0, 1 mM EDTA, 1.5% (v/v) SDS with 0.8 mg/mL proteinase K (Qiagen) and incubated at 56 °C for 1 hr. Deproteinated DNA was extracted with a Qiagen PCR purification kit according to the manufacturer's instruction. Extracted DNA was separated on a 3.5% NuSieve GTG agarose gel (Lonza) and the di-nucleosome band (~300 bp) isolated by gel-excision. Excised DNA was purified using the Nucleospin Gel and PCR cleanup kit (Macherey-Nagel) and divided into two equal pools. One pool was heat denatured at 95 °C for 5 min and snap cooled at 4 °C for 5 min using a thermocycler. DNA samples were separated on a 1% agarose gel at 100 V for 45 min with ethidium bromide. Gel images were captured with an Alpha Innotech Fluorchem 5500 and band intensity quantified with FIJI/ImageJ (1.53c), and images processed in Adobe Photoshop (22.3.2). Crosslinking efficiency was determined by normalising the native/crosslinked DNA band in the heat denatured samples, to their respective non-denatured samples.

Statistical analysis

Animals in each sample group were randomly selected. Sample group size (n) is indicated in each figure legend, while all experiments were carried out in at least three biological replicates. Statistical analyses were performed with Prism 4.0 software. The method used for each analysis is indicated in the corresponding section.

Resource sharing

*P. waltl* PacBio subreads (fastq and fasta files) are available under BioProject PRJNA353981 (SRR8594170).

Curated telomeric *A. mexicanum* contigs:
AMEXG_0030015054
AMEXG_0030017464
AMEXG_0030055159
AMEXG_0030100355
AMEXG_0030103279

Curated *P. waltl* PacBio subreads:
m150603_101004_42203_c100824942550000001823179511031565_s1_p0/101111/0_9897
m150807_080157_42237_c100857002550000001823190201241635_s1_p0/134571/0_11884
m150807_200551_42203_c100857072550000001823190201241664_s1_p0/148381/623_14254
m150603_101004_42203_c100824942550000001823179511031565_s1_p0/153040/0_14744
m150530_031530_42237_c100824912550000001823179511031592_s1_p0/19016/0_11208
m150722_021945_42237_c100854372550000001823189501241651_s1_p0/30177/10554_3509
2
m150710_220507_42237_c100834192550000001823181811251501_s1_p0/31594/0_13353
m150731_195104_42203_c100860772550000001823190601241674_s1_p0/49492/3359_19022
m150810_020230_42203_c100857222550000001823190201241650_s1_p0/811/0_11237
m150722_151724_42237_c100854372550000001823189501241654_s1_p0/93902/4596_16756

Illumina reads used for example telomeric region in *P. waltl*:
gnl|SRA|SRR6001099.138434328.1
gnl|SRA|SRR6001099.152158295.1
gnl|SRA|SRR6001099.209958579.1

**Methods references**

31. P. Ferretti, J. P. Brockes, Culture of newt cells from different tissues and their expression of a regeneration-associated antigen. *J Exp Zool* **247**, 77-91 (1988).
32. A. Joven, M. Kirkham, A. Simon, Husbandry of Spanish ribbed newts (Pleurodeles waltl). *Methods Mol Biol* **1290**, 47-70 (2015).
33. P. W. Tank, B. M. Carlson, T. G. Connelly, A staging system for forelimb regeneration in the axolotl, Ambystoma mexicanum. *J Morphol* **150**, 117-128 (1976).
34. N. P. Bordzilovskaya, Dettlaff, T.A., Duhon, S.T., Malacinski, G.M., *Developmental-stage series of axolotl embryos*. Developmental Biology of the Axolotl. Oxford University Press, New York, (1989).
35. M. H. Yun, P. B. Gates, J. P. Brockes, Sustained ERK activation underlies reprogramming in regeneration-competent salamander cells and distinguishes them from their mammalian counterparts. *Stem Cell Reports* **3**, 15-23 (2014).
36. N. J. O'Callaghan, M. Fenech, A quantitative PCR method for measuring absolute telomere length. *Biol Proced Online* **13**, 3 (2011).

37. J. D. Henson *et al.*, The C-Circle Assay for alternative-lengthening-of-telomeres activity. *Methods* **114**, 74-84 (2017).
38. M. R. Corces *et al.*, An improved ATAC-seq protocol reduces background and enables interrogation of frozen tissues. *Nature Methods* **14**, 959-962 (2017).
39. A. J. Cesare, C. M. Heaphy, R. J. O'Sullivan, Visualization of Telomere Integrity and Function In Vitro and In Vivo Using Immunofluorescence Techniques. *Current Protocols in Cytometry* **73**, 12.40.11-12.40.31 (2015).

**Supplementary Text**

Telomeric repeat frequencies

The frequency of each of the possible 48 hexamers ending in GGG and beginning with A, C or T (to prevent counting overlapping hexamers) were counted per read in 1 kb bins. The "telomeric region" of a sequence was defined as a region where the frequency of TTAGGG reached its peak. A "sub-telomeric region" of a sequence was defined as a region where the frequency of TTAGGG begins to increase before reaching its peak. The sequencing error rate inherent in PacBio reads (~15%) does not impact this analysis since the errors are evenly distributed across the length of the read. Moreover, the presence of telomere variants in *P. waltl* was confirmed using Illumina reads and fluorescent in situ hybridization.

TERT detection in newt genome

In the axolotl genome, *Tert* is flanked by *Clhc*1 (upstream) and *Top*1.2 and *Mylcd* (downstream). A variety of BLAST iterations did not detect TERT in the *P. waltl* genome, although *Clhc*1 and *Mylcd* were identified. Due to the fragmented nature of the current *P. waltl* genome assembly, one may argue that *Tert* exists in the genome but is misassembled. Importantly, the *P. waltl* transcriptome is exhaustive, and querying for the human, *Xenopus,* or axolotl *Tert* gene sequence within the *P. waltl* transcriptome does not retrieve any significant hits. Further, Blast iterations do not detect *Tert* within the PacBio long-read whole-genome dataset generated in this study. We therefore conclude that newt *Tert* has diverged beyond detection by homology.

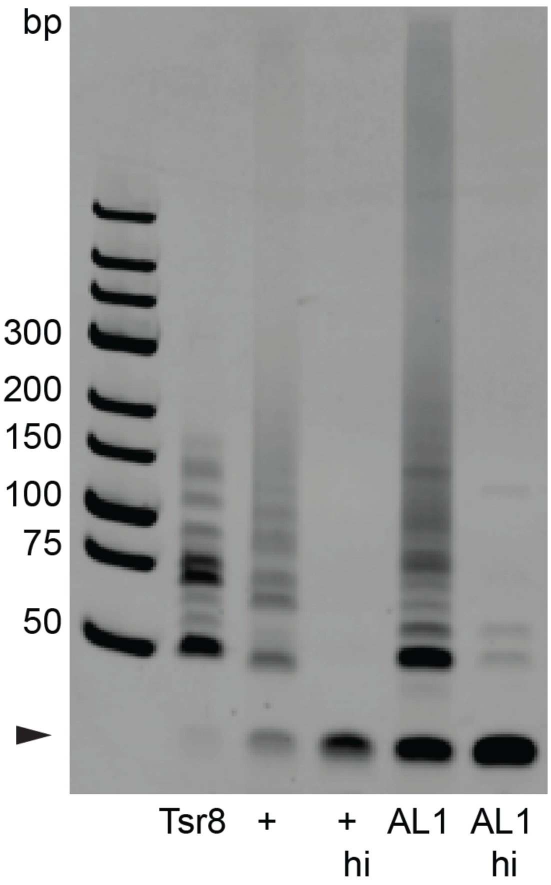


**Fig. S1. Telomerase activity in *A. mexicanum* extracts.** Telomerase enzymatic activity determined by TRAP assay in *A. mexicanum* AL1 cell extracts. TRAPeze telomerase positive human cell extract (+) is used as positive control. Tsr8=telomere consensus template (additional positive control). hi=heat inactivation (negative control). A 36-bp internal control for amplification efficiency was run for each reaction (arrowhead, n=4).

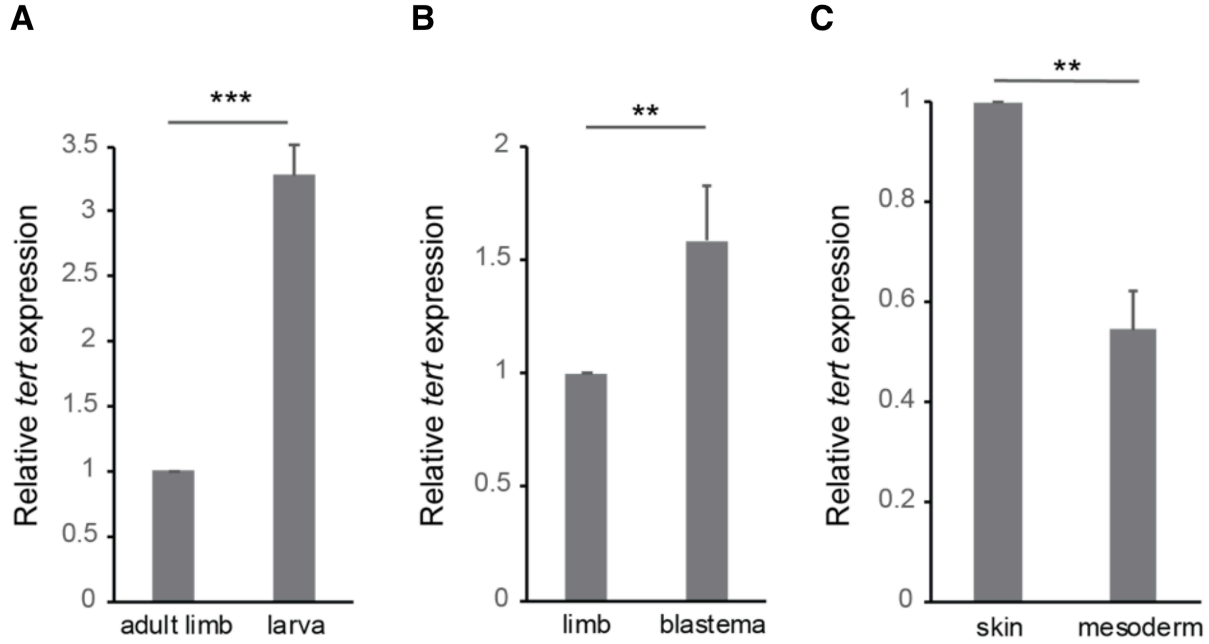


**Fig. S2. *Tert* expression in *A. mexicanum* tissues during development, regeneration and homeostasis.** (A to C) qRT-PCR analysis of axolotl *Tert* expression levels in the indicated tissues. Gene expression levels were normalised to those of *ef1*-α. Values represent the mean ± SD of at least three independent experiments. **p<0.01, ***p<0.001 (Student's t test).

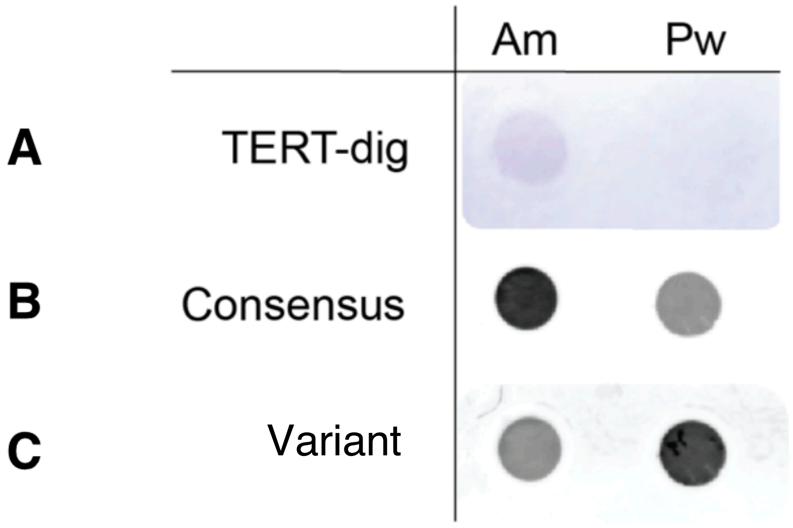


**Fig. S3. Dot blot analysis of TERT in *A. mexicanum* and *P. waltl* genomes**. *A. mexicanum* and *P. waltl* genomic DNA were hybridized at low stringency conditions with a TERT-digoxigenin probe (A) and developed using immunological detection (purple signal). The same membrane was then stripped and re-probed with either consensus (B) or variant (C) telomere probes as loading controls, as indicated. Data are representative of n=3 biological replicates.

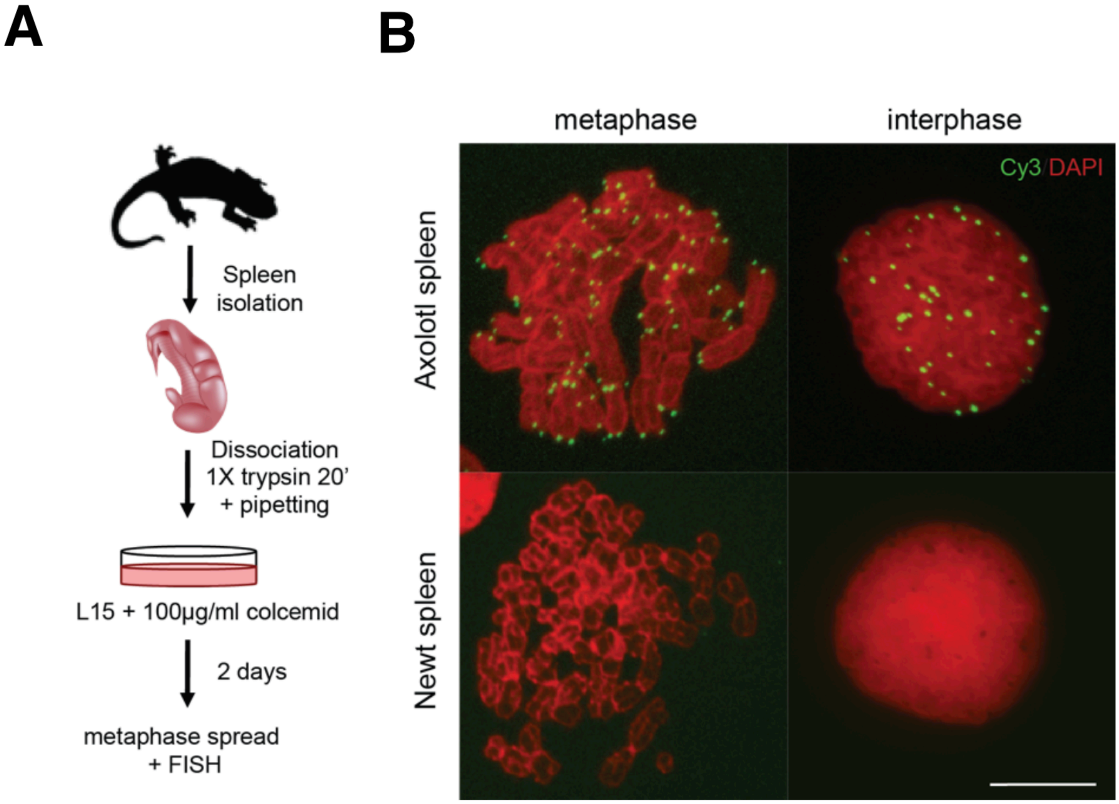


**Fig. S4. Canonical telomere signals are absent in spleen cultures derived from the newt *N. viridescens*.** (**A**) Schematic of the experimental procedure. (**B**) FISH assay using a Cy3-labelled telomere consensus probe on interphase nuclei or metaphase spreads derived from axolotl and newt spleen cultures. No signal was detected in *N. viridescens* cells (n=3). Scale bar: 20µm.

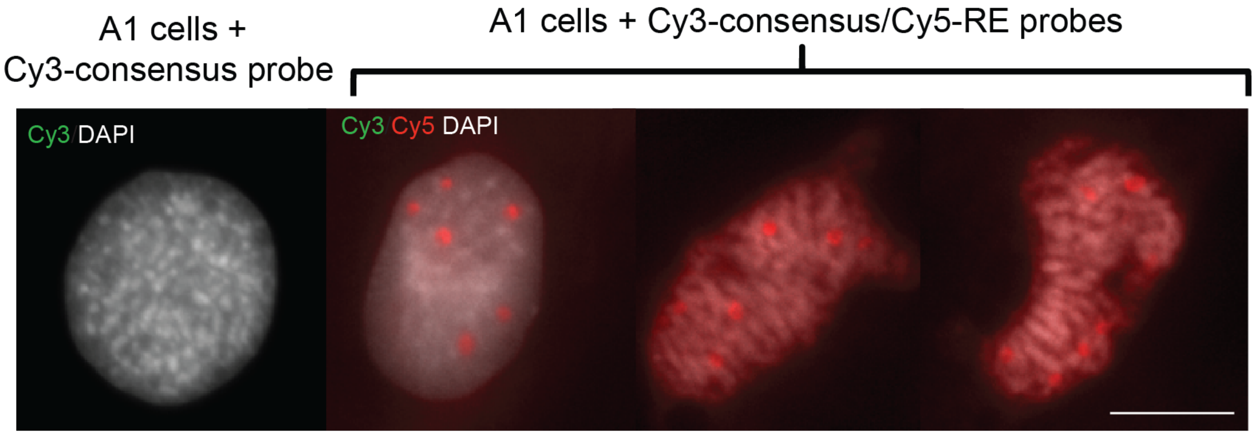


**Fig. S5. Fluorescence in situ hybridization of a common repetitive element in *N. viridescens.*** FISH assay using a Cy3-labelled telomere consensus probe with/without a Cy5-labelled probe against a common repetitive element (RE, consisting of consecutive repeats of the sequence GGTATAGGTATA) on newt A1 cells. Note positive signal for the Cy5-RE probe (n=3). Scale bar: 20µm.

A

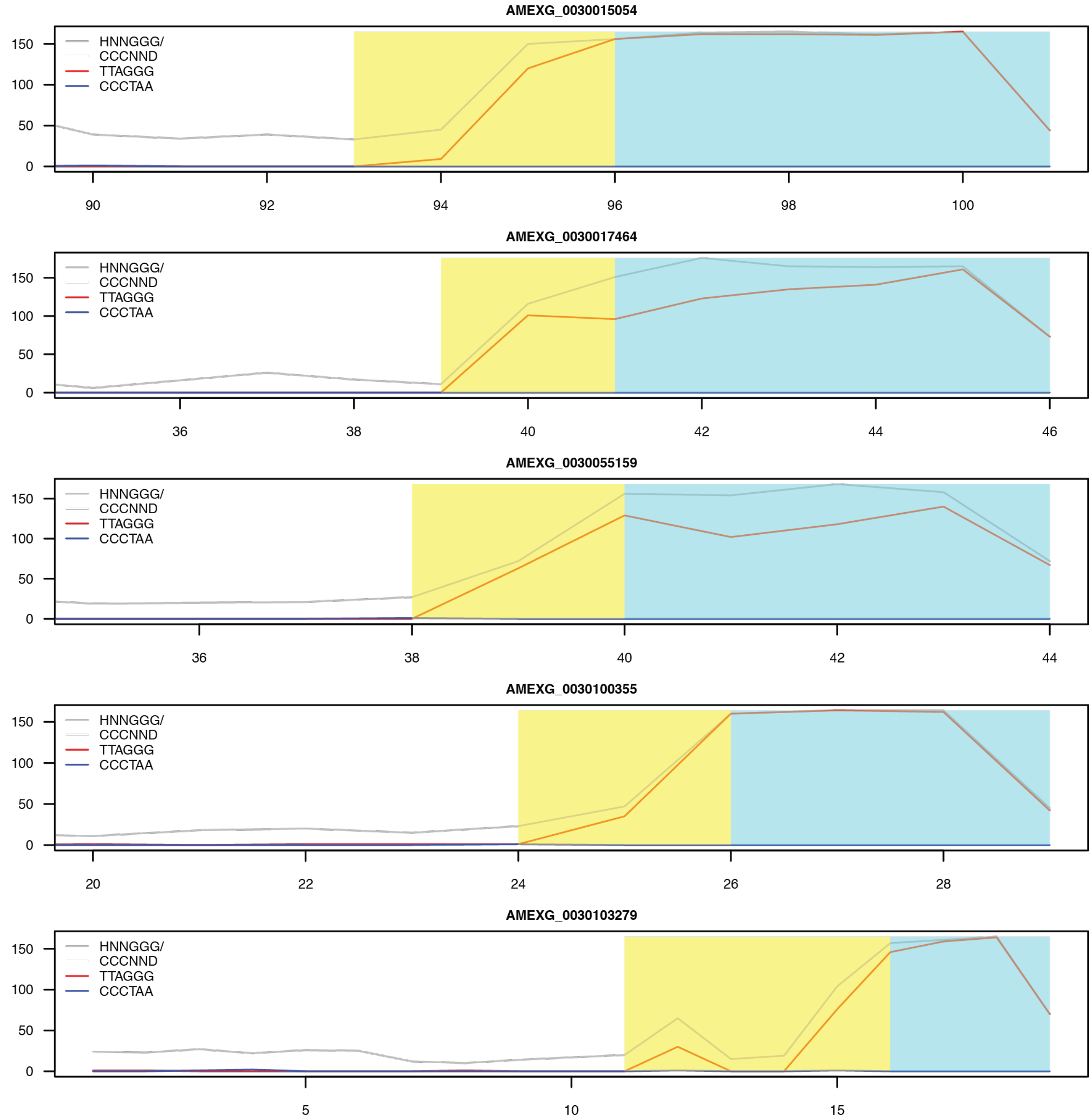
AMEXG_0030015054
HNNGGG/
CCCNND
TTAGGG
CCCTAA
0
50
100
150
90
92
94
96
98
100
AMEXG_0030017464
HNNGGG/
CCCNND
TTAGGG
CCCTAA
0
50
100
150
36
38
40
42
44
46
AMEXG_0030055159
HNNGGG/
CCCNND
TTAGGG
CCCTAA
0
50
100
150
36
38
40
42
44
AMEXG_0030100355
HNNGGG/
CCCNND
TTAGGG
CCCTAA
0
50
100
150
20
22
24
26
28
AMEXG_0030103279
HNNGGG/
CCCNND
TTAGGG
CCCTAA
0
50
100
150
5
10
15

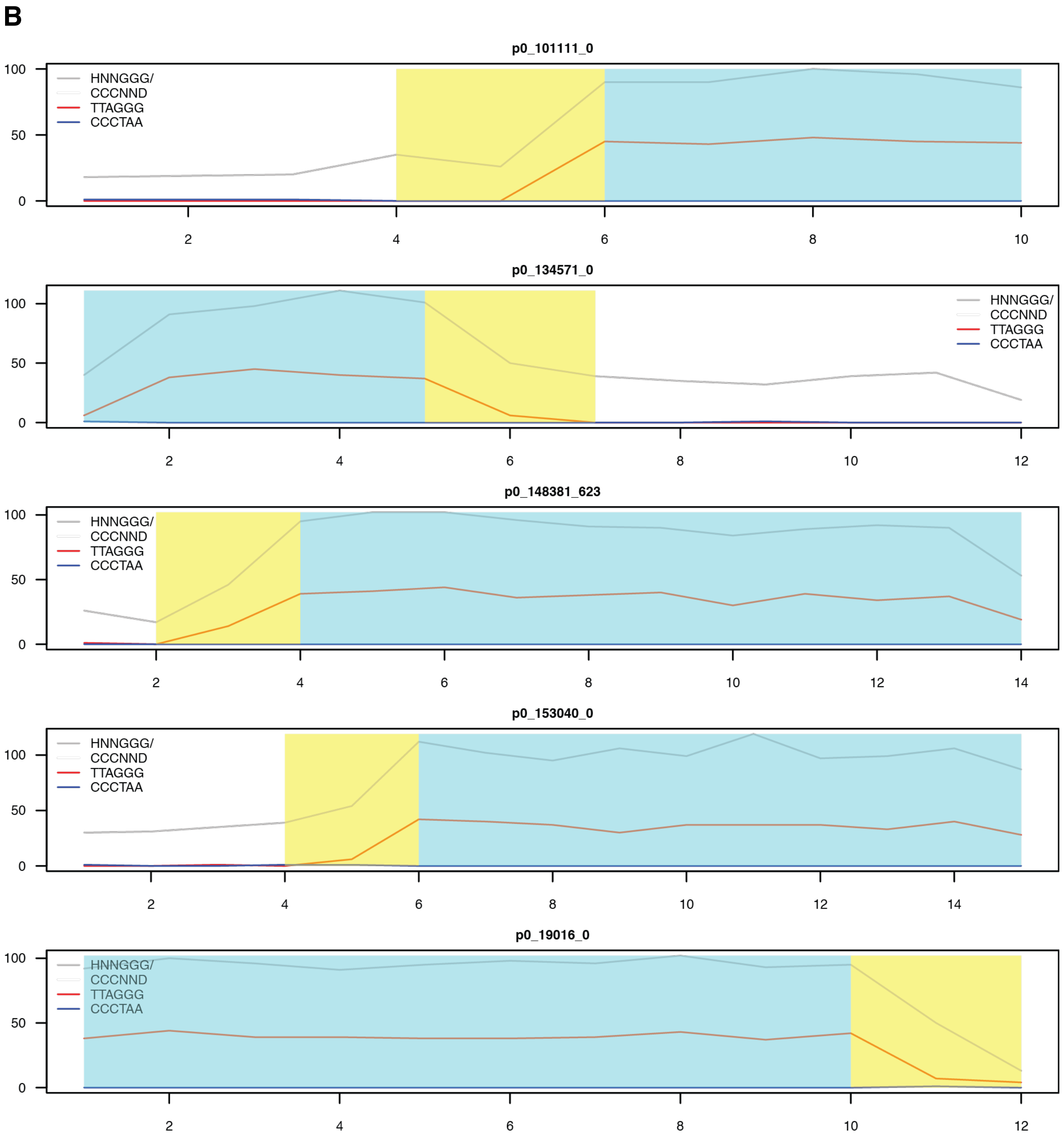


**Fig. S6. Representative *A. mexicanum* (A) and *P. waltl* (B) telomeric PacBio genome sequencing reads.** Sub-telomeric (yellow) and telomeric (blue) regions are highlighted. Counts of total telomeric variants (HNNGGG, grey), telomere consensus hexamer (TTAGGG, red) and its complement (CCCTAA, blue) along each read are shown.

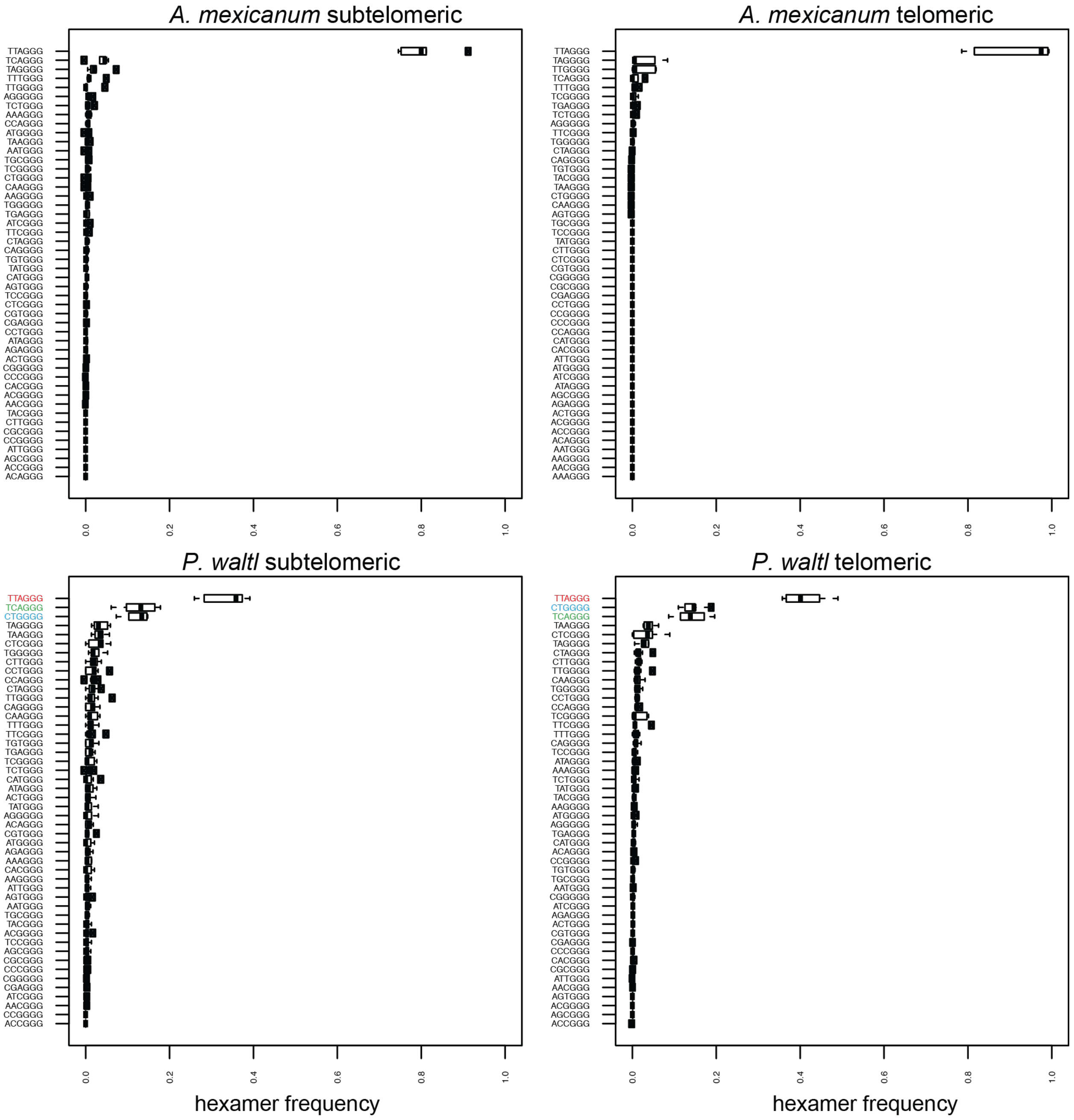


**Fig. S7. Frequency of HNNGGG hexamers in sub-telomeric and telomeric regions of *A. mexicanum* and *P. waltl* curated telomeric reads from PacBio genomic data.** The three most prevalent hexamers in *P. waltl* (bottom) are coloured according to Fig.2. Note that the variant probe used in this study is composed of the three most frequent hexamers (coloured).

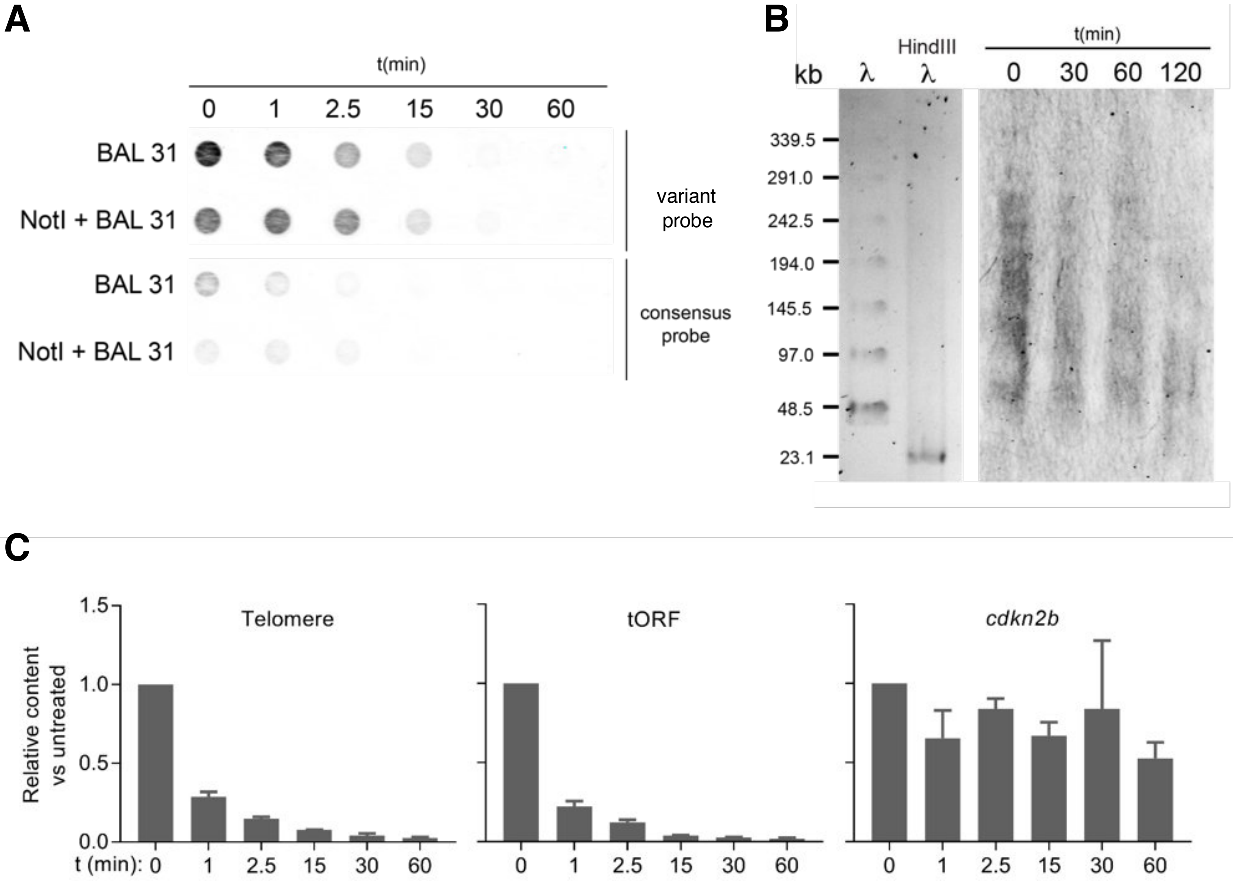


**Fig. S8. The variant probe binds specifically to *P.waltl* chromosome termini.** (**A**) Dot blot of *P. waltl* DNA digested with BAL-31 exonuclease ± NotI to expose the non-telomeric regions to BAL-31 digestion for the indicated times and hybridized with variant or consensus probe. (**B**) Pulse field gel electrophoresis (PFGE) and Southern blotting of *P. waltl* DNA digested first with BAL-31 exonuclease, then rendered as telomere restriction fragments. The blot was hybridized with variant probe and shows a BAL-31-dependent decrease in telomeric signal consistent with terminal localization of the variant repeats. (**C**) *P. waltl* genomic DNA was treated with BAL-31 exonuclease for the indicated times. Thereafter, telomeric, tORF (telomeric open reading frame, discussed in Fig. 4E), and genomic *cdkn2b* (non-telomeric control) content was measured using qPCR. Data are expressed as ratio of target content to untreated (-BAL-31) samples and normalized against genomic *ef1α* using the ΔΔCT method. Data represent mean ± SD of two technical replicates. (*p<0.05, **p<0.01, ****p<0.0001, Student's t test). Note that telomere and telomeric ORF (tORF) content decrease with exposure to BAL-31 as expected, indicating their localization at chromosomal ends, whereas a non-telomeric sequence (*cdkn2b*) is unaffected by exonuclease treatment.

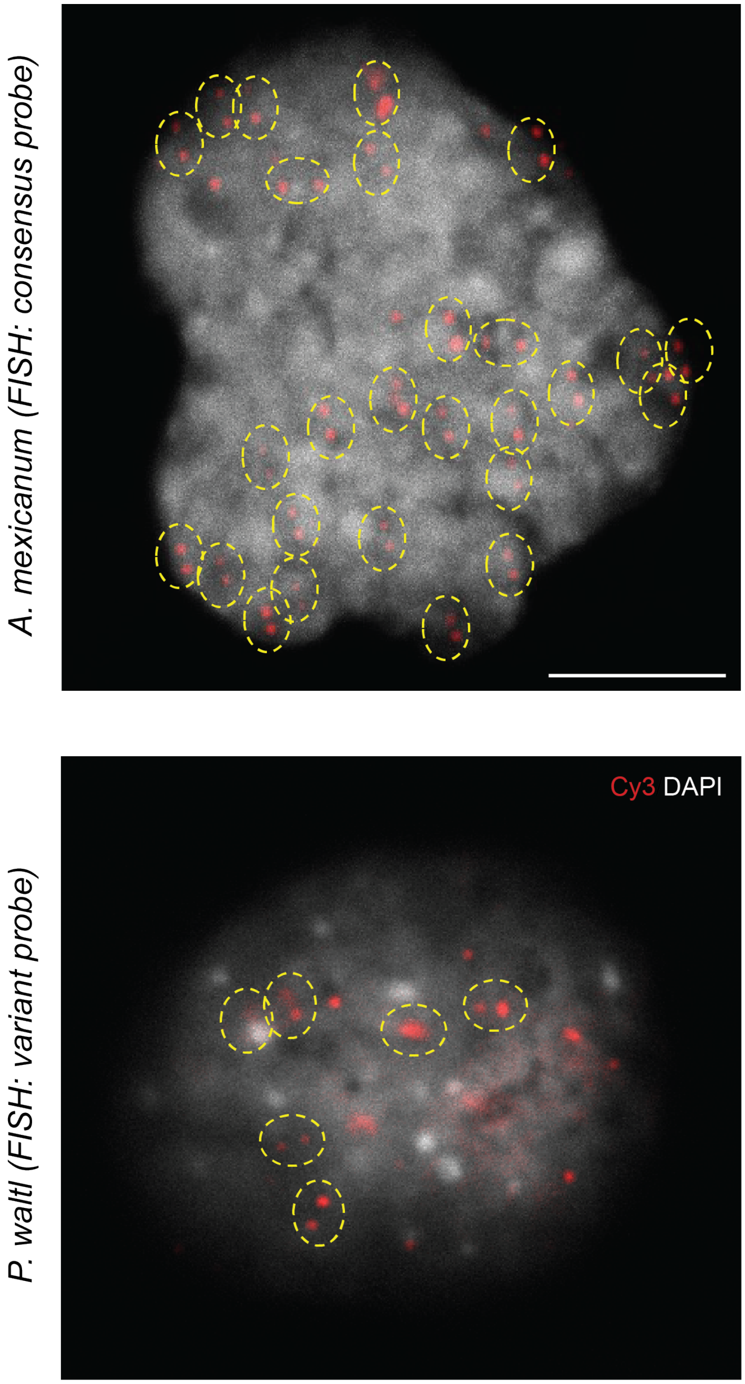


**Fig. S9. FISH with the variant probe reveals punctate nuclear foci consistent with telomeres in *P.waltl* nuclei**. Magnification of Fig. 2E showing FISH labelling of punctate nuclear foci consistent with telomeres using the consensus or variant probes in samples from *A. mexicanum* (top) and *P. waltl* (bottom) embryonic cells, respectively (yellow circles) Scale bar: 25µm.

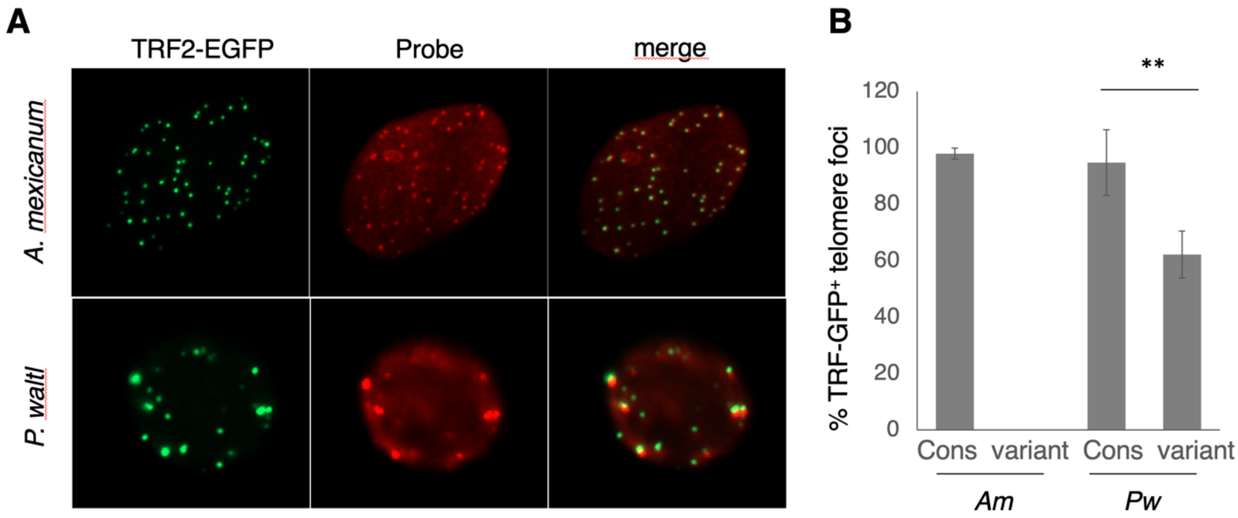


**Fig. S10. TRF2 colocalizes with telomeric sequences in axolotl and *P. waltl*.** (**A**) Representative images of *A. mexicanum* or *P. waltl* cells following electroporation with TRF2-EGFP-N2 and subsequent visualization of telomeres using the consensus probe through fluorescent in-situ hybridisation under low stringency conditions. (**B**) Quantification of the percentage of telomere foci, as indicated for each probe, co-localising with TRF2-EGFP fusion protein. No signal is detected in *A. mexicanum* cells using the variant probe. Data represent mean ± SEM of three experiments; ≥20 cells scored per replicate. $^{**}p < 0.01$ (Student's t test).

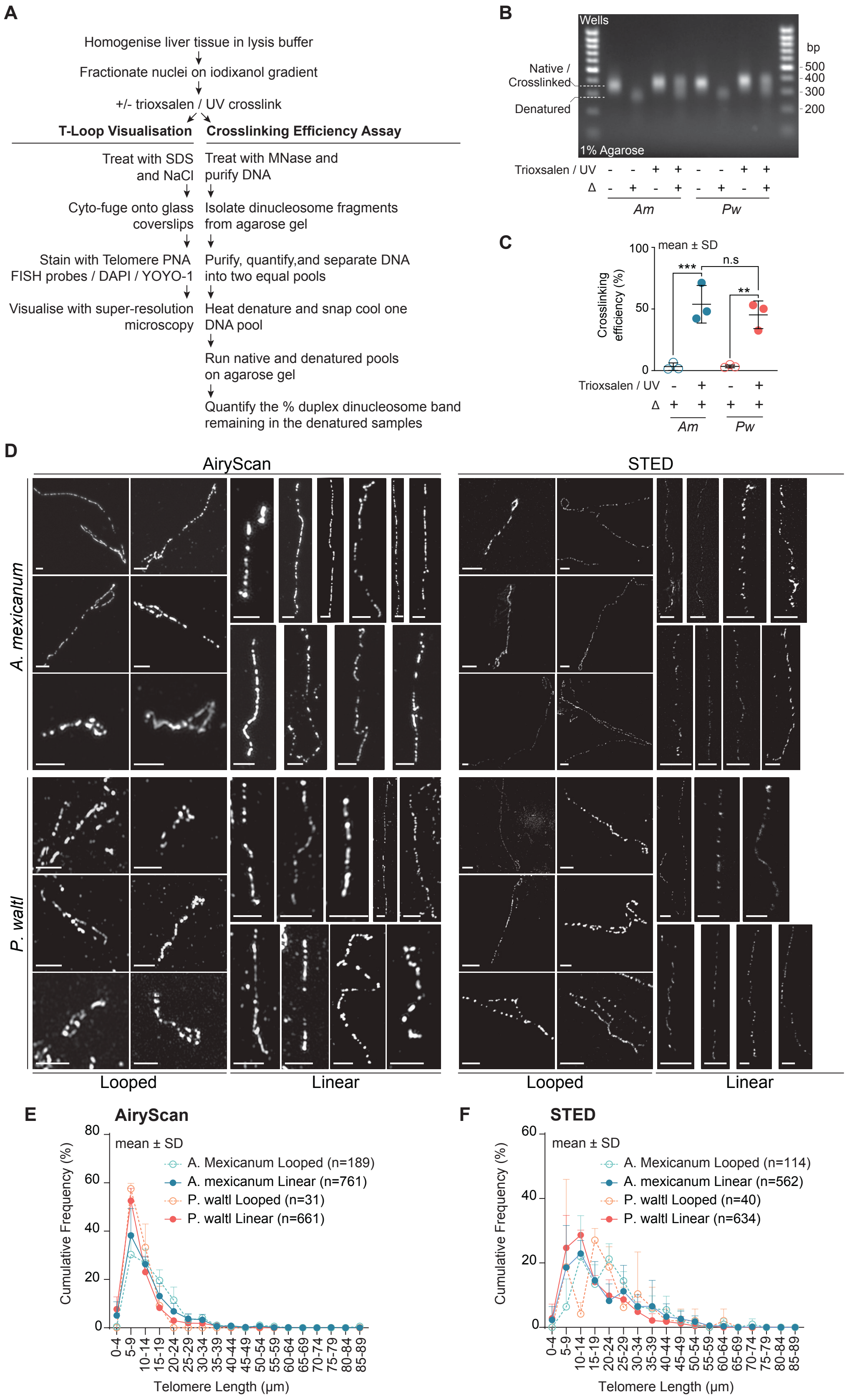
A
Homogenise liver tissue in lysis buffer
Fractionate nuclei on iodixanol gradient
+/- trioxsalen / UV crosslink
T-Loop Visualisation
Crosslinking Efficiency Assay
Treat with SDS and NaCl
Treat with MNase and purify DNA
Cyto-fuge onto glass coverslips
Isolate dinucleosome fragments from agarose gel
Stain with Telomere PNA FISH probes / DAPI / YOYO-1
Purify, quantify,and separate DNA into two equal pools
Visualise with super-resolution microscopy
Heat denature and snap cool one DNA pool
Run native and denatured pools on agarose gel
Quantify the % duplex dinucleosome band remaining in the denatured samples
B
Wells
Native / Crosslinked
Denatured
1% Agarose
bp
500
400
300
200
Trioxsalen / UV
Δ
Am
Pw
C
mean ± SD
Crosslinking efficiency (%)
100
50
0
***
n.s
**
Trioxsalen / UV
Δ
Am
Pw
D
AiryScan
STED
A. mexicanum
P. waltl
Looped
Linear
Looped
Linear
E
AiryScan
mean ± SD
Cumulative Frequency (%)
A. Mexicanum Looped (n=189)
A. mexicanum Linear (n=761)
P. waltl Looped (n=31)
P. waltl Linear (n=661)
Telomere Length (µm)
F
STED
mean ± SD
Cumulative Frequency (%)
A. Mexicanum Looped (n=114)
A. mexicanum Linear (n=562)
P. waltl Looped (n=40)
P. waltl Linear (n=634)
Telomere Length (µm)

**Fig. S11. Sample preparation and super-resolution imaging of telomere macromolecular structure.** (**A**) Schematic of the sample preparation used in (B-D) below and Figure 3C-D. (**B**) Representative agarose gel used to determine crosslinking efficiency in *A. mexicanum* and *P. waltl* liver tissue. (**C**) Quantification of the crosslinking efficiency assay (n=3 biological replicates, one-way ANOVA Tukey's multiple comparisons, ** $p < 0.01$, *** $p < 0.001$). (**D**) Representative looped and linear telomeres in *A. mexicanum* and *P. waltl* liver tissue from AiryScan and STED microscopy analysis (scale bars indicate 2 μm, n=3 biological replicates). (**E**) Histogram of total telomere length (loop + tail) in *A. mexicanum* and *P. waltl* samples captured by AiryScan and (**F**) STED microscopy (n=3 biological replicates).

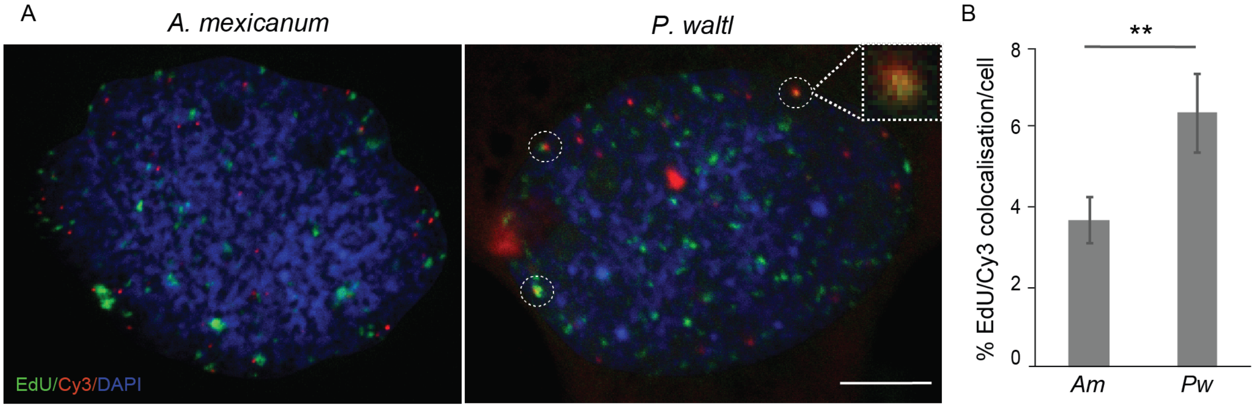


**Fig. S12. EdU is incorporated at *P. waltl* telomeres outside of S-phase.** (**A**) Representative images of telomere (Cy3-labelled consensus (*Am*) or variant (*Pw*) probes) colocalization with punctate EdU foci in *A. mexicanum* or *P. waltl* cells. S-phase cells incorporate EdU diffusely throughout the nucleus and were excluded from subsequent analysis. (**B**) Quantification of EdU/Cy3 colocalization in non-S-phase cells. Data represent mean ± SEM of three experiments; ≥ 20 cells scored per replicate. $**p < 0.01$. Scale bar: 20µm.

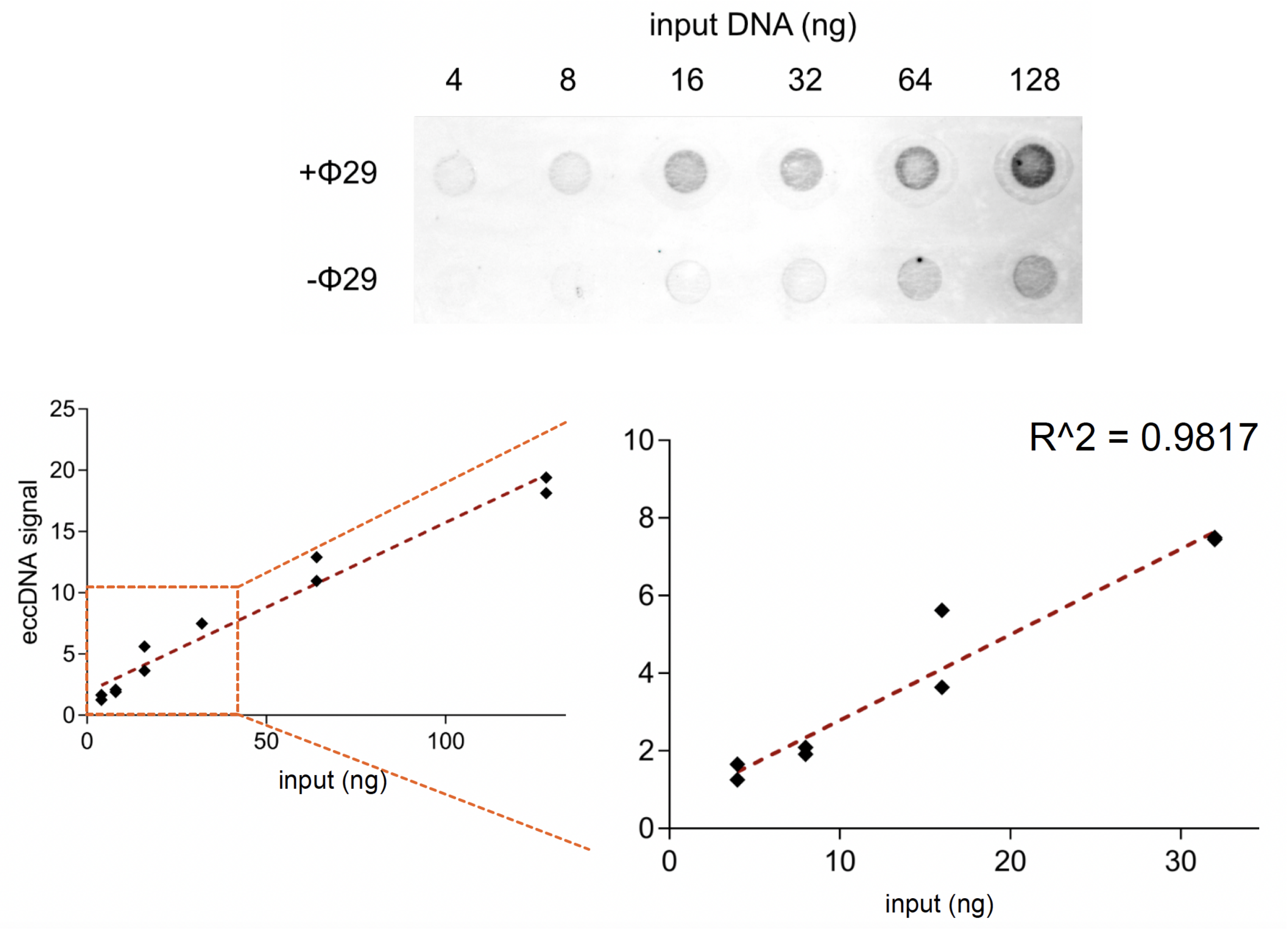


**Fig. S13. Rolling circle amplification assay for the detection of telomere circles in newt tissues.** Representative RCA assay with increasing amount of P. waltl gDNA from testes (top); dot blots were hybridised with variant probe as described (Methods). Bottom: Determination of the assay's linear range. The assay is linear between 4 and 32ng gDNA. Note that 10ng are used as input in all our described experiments, thus they fall within the linear range.

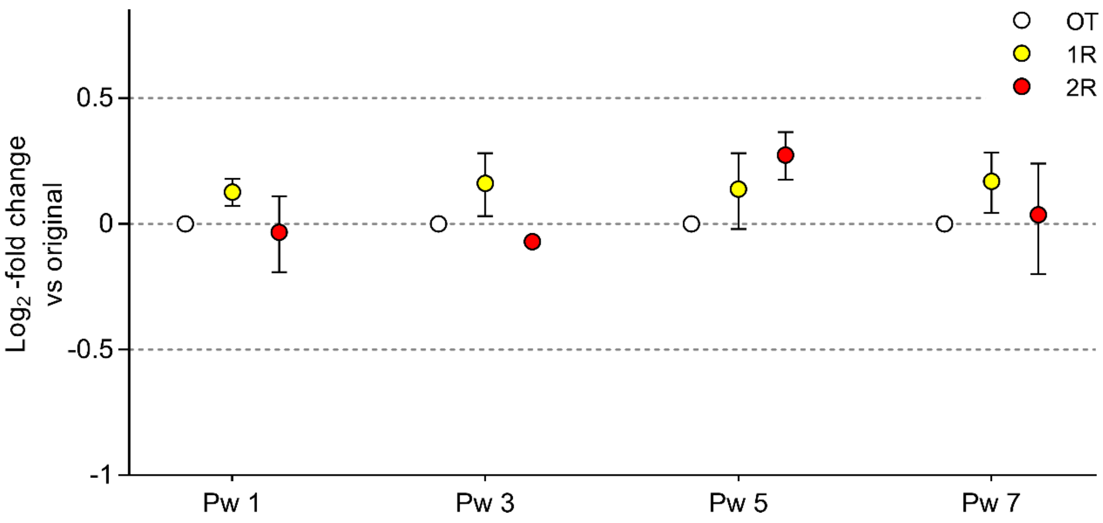


**Fig. S14. Serial regeneration rounds do not alter the copy number of a non-telomeric sequence.** Changes in the copy number of the genomic *ef1α*, assessed by qPCR throughout multiple rounds of regeneration in *P. waltl* individuals. The samples used in this experiment -and their nomenclature- correspond with Fig. 4F. Results are expressed as fold-change in *ef1α* copy number of first- (1R) and second-regenerated (2R) tails compared with the original (OT) and normalized to *cdkn2b* using the $^{\Delta\Delta}$CT method. Data represent mean ± SEM of four technical replicates. No statistically significant changes in relative content are detected across multiple regeneration rounds (one-way repeated measure ANOVA followed by Tukey's *post hoc* test).

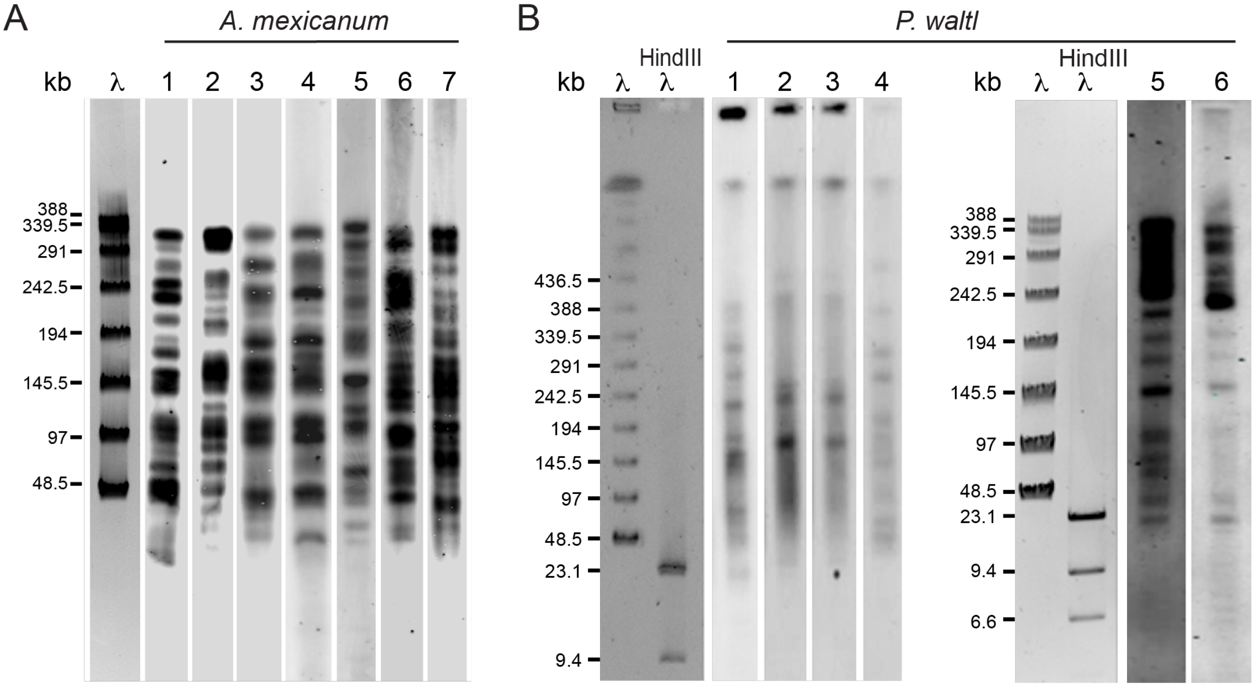


**Fig. S15. Inter-individual variability in telomere banding patterns in *A. mexicanum* and *P. waltl.*** Telomere restriction fragment analysis by pulse field gel electrophoresis and Southern blotting of DNA isolated from 7 different *A. mexicanum* (**A**) and 6 different adult *P. waltl* (**B**) individuals of the same age, hybridized with telomere consensus (A) or variant (B) probes.

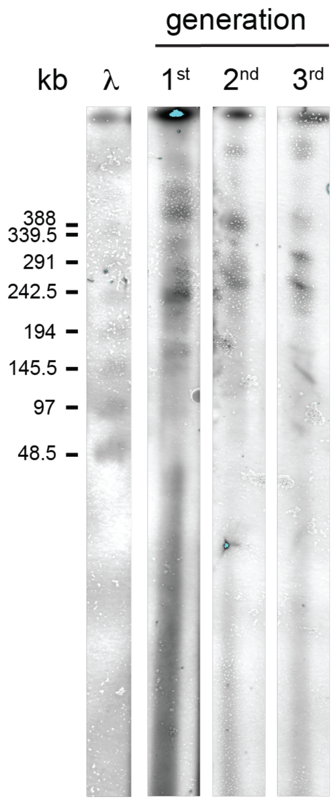


**Fig. S16. Telomeres do not shorten in *P. waltl* over three generations.** Representative telomere restriction fragment analysis by pulse field gel electrophoresis and Southern blotting of genomic DNA from *P. waltl* newts belonging to 3 consecutive generations, hybridized with the variant telomere probe.

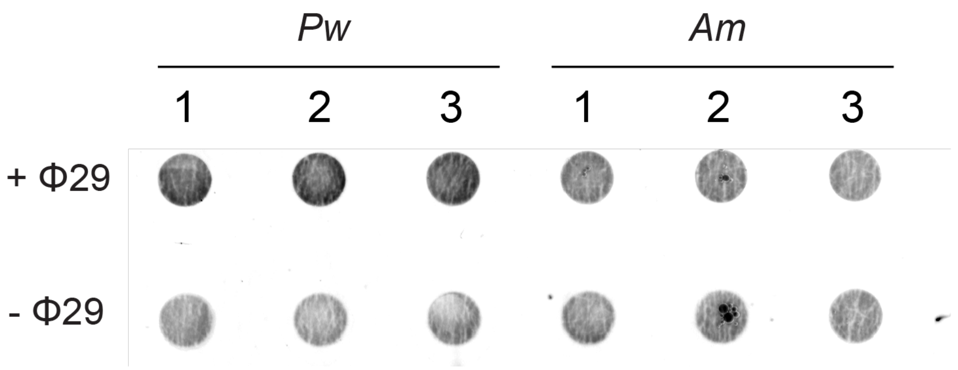


**Fig. S17. Rolling circle amplification assay in salamander testes samples.** RCA assay of *P. waltl* and *A. mexicanum* samples from testes from three individuals, run in a single membrane. Dot blots were hybridized with the variant probe. Blot corresponds to uncropped image of Fig. 4I.

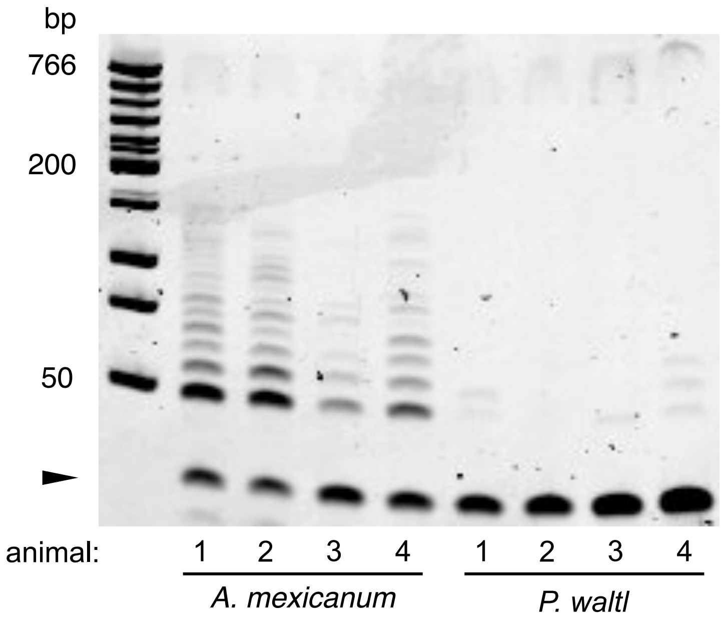


**Fig. S18. Telomerase activity is not detected in the *P. waltl* germline.** Telomerase enzymatic activity determined by TRAP assay in *A. mexicanum* and *P. waltl* testes extracts (n=4). A 36-base pair internal control for amplification efficiency was run for each reaction (arrowhead).

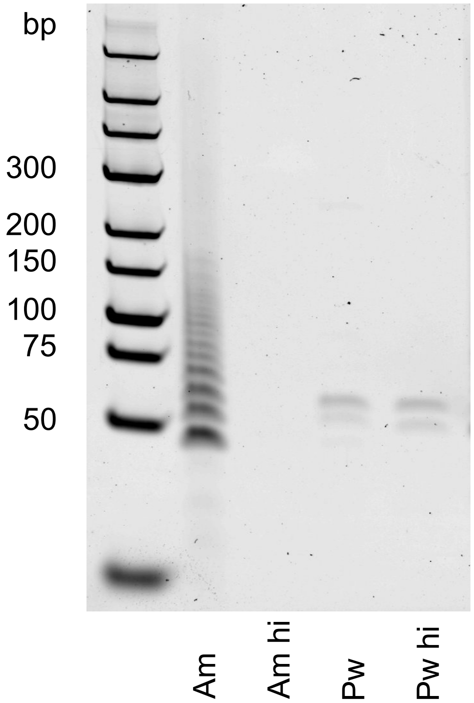


**Fig. S19. Telomerase activity is not detected in *P. waltl* TRAP assays when amplifying with variant telomere sequence.** Telomerase enzymatic activity as determined by TRAP assay in *A. mexicanum* and *P. waltl* testes extracts using the variant telomere sequence for product amplification in the PCR step. hi=heat inactivation (negative control). A banding pattern resulting from telomerase activity is present in the *Am* extract because the variant primers anneal to consensus telomerase extension products (n=4).

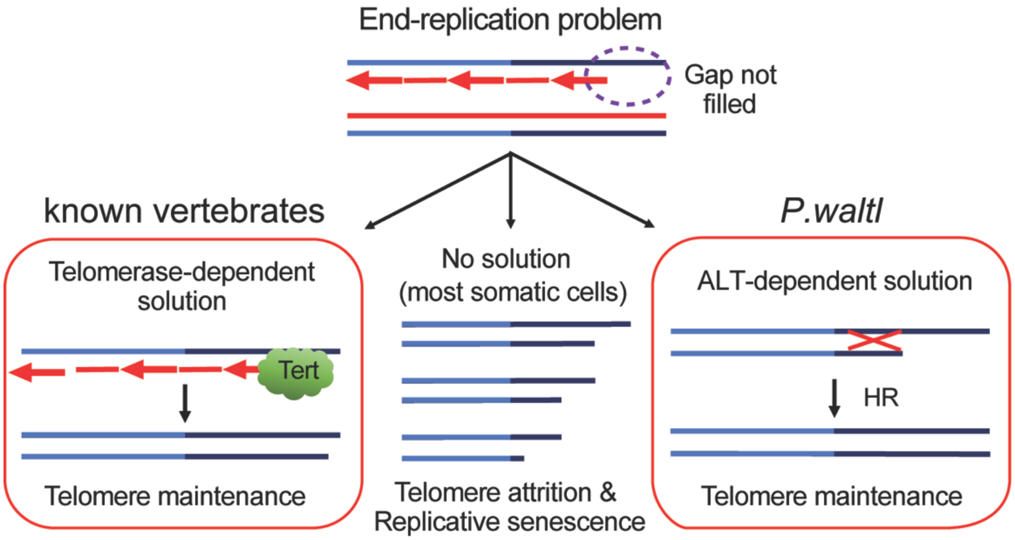


**Fig. S20. *P. waltl* employs the ALT pathway to maintain telomere length throughout lifespan.** Schematic model for how replicative cells overcome the end-replication problem in known vertebrates. *P. waltl* constitutes an exception to the rule of telomerase-dependency. HR: homologous recombination.

**Table S1. Protein sequences of genes flanking the *Tert* locus of *A. mexicanum*.**

| ***Am*** **gene** | protein sequence |
|---|---|
| Top1.2 | MMNSAFDGKDIETARGSTKSLSSSEKTDQSKENKKENLNDGIKKTSNTSDRKSKSLSKGS<br>SRKDADGTLTDEVFEGKTPESSQGSKRSLSSTNPNENQGEKLNRSVTEKNLKHEKRKSKS<br>STKGLKQTDNLDGSLMKEEYIEKDLEFYPGPTKSCRNAVDDDEDKENREELINVGVKEK<br>RSKQDEREKRKSKSSSKDLKQKDILDGKSLNYTSSGKGIQSRTMDIQEKKSKDGETMGS<br>KSKQDVDLNSEHEVNHFKENQNEKRKSRTKSIGSKKDDYRPKPFKEEPEDSDFLTIHNER<br>KRTQLTVQDGEAKEVAGTAMKEVLKQKKLETPSRSKKGLEEEKKVNQDHDKINGNQEK<br>SNKKEKRKSKIKSMDLKQIDFDGGESIKGELDDSVFLPAPMEGKREKRPRSEDTDGTPGS<br>KTFKKEPEDQDFVIIPKDVKRKRSMAEGAEEALSERKIKKIKTEINDQELSTKEEKSEQEIQ<br>LKKKKKKKKEEKEVEIKPKKEKKEGNQETKKKKTKEERELIADIKAKKKEEEEKQQWR<br>WWEEQKYEDGINWMFLEHKGPVFAPLYEPLPENVNFYYDGKPLKLCKAAEEVTSFFAK<br>MLDHEYTTNDVFRKNFFMDWRKEMTLEEKIIIADLGKCDFTEMFTYYKGKMEEKRNPT<br>KEQKLANKEANEKVIQEYGFCIMDHHRERIGNFKIEPPGLFRGRGDHPKQGMLKKRILPE<br>DVIINCSKDSKIPEPPEGHKWKEVRHDNTVTWLASWTENVQGSNKYVMLNANSRLKGE<br>KDWEKYEVARKLKTCVDRIRAQYHEDLKSKEMKLRQRAVALYFIDKLALRAGNEKEEG<br>ETADTVGCCSLRVEHIKLYPELDGEKHVVEFDFLGKDCIRYYNKVPVEKQVFKNLQLFM<br>EEKQGGDDLFDRLNTPMLNKHLQSLMEGLTAKVFRTYNASITLQDQLKKLTDPDEPVPS<br>KILSYNRANRAVAILCNHQRAAPKTFEQSMANLQTKIDAKKEKISLAKSELKQAKLDAK<br>SSDSEKLKKVLESKKKAVKRAEEQLMKLEVQATDREENKQIALGTSKLNYLDPRISVTW<br>CQKWGVGLEKIYNKTLRDKFAWAINMSDASFVFK |
| Clch1 | MSALDVSLNKSPALPPITSQCDRAFLKSIDKYIAEELRQLGCDGERNIENIYLIYRAVFDKV<br>IEYLTSYKNILSAVKQEYDTLIDSITLGQKDAFYLNEKLKSVTIDSNRLIYYKRRATQLQD<br>KIEIIEADSAKYFNLIEDKKAAKKCTTTCSTEPESNISSVQINPTIRMPGMTINESLDIDALA<br>KYHFQLEKKMLCLKTEMKTKYISAKMKAELDQSLELALLARDEAESINTSLKLSCKKRK<br>LIVSAIRTWATSDKRKTLSEFLYERIKTEDSSEDEQVAENVFGDEDPSNIKEGDDLLEYIER<br>FNELFEAGQYESASVHAANCPREILRNMHTMERFKAAPFVEGKVPPLMFFFEALITSSSTT<br>KHPVSACMTLEAIKCALMQKRLDLVTHWITQQRLTFSEALGDAIFEYFQTEHHKRAECL<br>ALALAQIIYRQCGVHRKTALCMCLESQICGAVEFIHQCKKFSLDDLLFLLKKCPTNELIEN<br>LTREWNGKKAIVSTGQAVLLLIDADHKEIGFQMLEHFDTHGTCTLEDVILSDEVCTIEGW<br>TEIADQCSRNDYIPLSQKIKSVLTSQDGIVEMSPDDYDALLMEHVFL |
| Rps27a | MQIFVKTLTGKTITLEVEPSDTIENVKAKIQDKEGIPPDQQRLIFAGKQLEDGRTLSDYNIQ<br>KESTLHLVLRLRGGAKKRKKKSYTTPKKNKHKRKKVKLAVLKYYKVDENGKIHRLRRE<br>CPSEECGAGVFMASHFDRHYCGKCCLTYCFNKPEDK |
| Mtif2.L | KMNRAMLQMLETLVQFTTACRQLRSLHSGKAQRMHWRSTQCSVYPKCASQQHQPLCQ<br>TDTVLNVSLTHCRLLATKEGKSPKKYKVFPKKPNLDKQEVEIWNKMTVEDLARAMDKD<br>IDHVYEALLHTTLNLDELEPQSVLDEVWIKEAVKKSGMKFKYAKLKEEKLRENKDAVN<br>RPPAEPALLTPRPPVVTIMGHVDHGKTTLLDKLRETQVAAMEAGGHHTAHWGFMEKQH<br>YLTSYERPRWQQWRLAGITQHIGAFNVHLSSGEKITFLDTPGHAAFSAIRARGAHVTDIVI<br>LVVAAEDGVMKQTVESIRHAKEAKVPIILAVNKCDKPEADPERVKKELLAHNVVCEEFG<br>GDVQAIHISALKGDNLLALVEATVTLAEVLELKADATGPVEGVVVESRTDKGKGPVTTA<br>IIQRGTLKKGCVLVAGTSWAKVRLMFDENGTGMEAAGPSMPVEIVGWKELPSAGDEILQ<br>VDSEQRAEKLLNGENMWKSRRKSREVVEWRKYVEEQEKVKQDVDVIEAKQKEHRETH<br>KKQLEALGNLHWRQRKASMYRANKQVWGSRSKENVEDESLTLPVIIKGDVDGSVEALL<br>GLIDSYDASDQCQLDLVHFGVGDISENDIDLAEAFKGKVI |
| Mlycd | MRLLSLRRAGRLLGSLPTRRSLSMDELLHLCVPLLPPYEARGRSQPPPELQSKSFMGQYQ<br>ALDHTGRSGLLRTLAVDFGRDHSRVAELCAAVLQVQDRGAAAMLQAEDRLRHQLCPR<br>YRLLFAHLSTAPDGVKFLLDLRADLLQSLEPAASEGPELREMNGVLKSMFSEWFSVGFL<br>NLERITWQSPCELLQKISESEAVHPVRSWTDMKRRVGPYRRCYVFTHSAMPGEPLIILHV<br>ALTNKIASNIQAIVKEVSTLDVEDVDKITAAIFYSISLAQQGLQGVELGNYLIKRVVKQLK<br>AEFSHLKDFSSLSPIPGFSKWFLGLLTSLKKEVGRNELFTDSECKEISDIVGEPITDALRHLI<br>SSNEWMRSERLIQALEPPLMRLCAWYLYGEKHRGFALNPVANFHLQNGAVMWRLNW<br>MADTSPRGLTASCGMMVNYRYFLEDTSGNSERYLRTKHIEASEQVLNLVSQFQTNSKL-<br>MQKKNFGKAIDRIKIKKMVECFVFGATLIK |

**Table S2. Accession numbers of homologues of genes flanking the Tert locus of *A. mexicanum*.**

| ***A. mexicanum*** | ***P. waltl*** | ***N. viridescens*** |
|---|---|---|
| *Top*1.2 | TRINITY_DN2120226_c0_g6_i4 | n/a |
| *Clch*1 | TRINITY_DN2033801_c0_g1_i3 | Cluster-64891.0_comp838596_c0_seq1 |
| *Rps*27a | TRINITY_DN2169082_c0_g1_i5 | Cluster-73776.37423_TRINITY_DN1056889_c0_g1_i4 |
| *Mtif*2.L | TRINITY_DN2165355_c2_g2_i4 | Cluster-35212.4_comp95911_c0_seq3 |
| *Mlycd* | TRINITY_DN2074082_c0_g3_i2 | Cluster-16005.1_comp89234_c0_seq1 |

**Table S3. Primer sequences.** Am: *A. mexicanum*; Pw: *P. waltl*; Nm: *N. maculosus*; Hd: *H. dunii*; Amac: *A. macrodactylum*. (*) indicates primers that were used as normalisers for tORF analysis as well as telomere size determination. ($^{\#}$) indicates primers that were used for Bal-31 digest qRT-PCR analysis. Telomere consensus standard oligomer consisted on 84 repeats of the TTAGGG hexamer.

| | *Name* | *Sequence (5' – 3')* |
|---|---|---|
| *Tert qRT-PCR* | Am *tert* forward | AAGCCTGACCATCAGCCATA |
| | Am *tert* reverse | AACCTGTAAGCCTGCAGCAA |
| | Am *ef1α* forward | AACATCGTGGTCATCGGCCAT |
| | Am *ef1α* reverse | GGAGGTGCCAGTGATCATGTT |
| *Bal31 qRT-PCR analysis* | Pw telomere forward | TTAGGGTAACGGCTGGG |
| | Pw telomere reverse | CGCAGCCCTAACCGTGA |
| | Pw *e1fα* reverse | ACTGTGCGGTGCTGATTGTTG |
| | Pw *e1fα* forward | GCTGACTTCCTTGGTGATTTCTTC |
| *Telomere size determination* | telomere consensus forward | CGGTTTGTTTGGGTTTGGGTTTGGGTTTGGGTT TGGGTT |
| | telomere consensus reverse | GGCTTGCCTTACCCTTACCCTTACCCTTACCCT TACCCT |
| | Am *hoxd10* forward | TTGAGCTTCATCCTGCGGT |
| | Am *hoxd10* reverse | CAGTAAGAGTGTGAACCTCAC |
| | Am *hoxd10* standard oligomer | GATCAGTAAGAGTGTGAACCTCACTGACAGGC AGGTCAAGATCTGGTTTCAAAACCGCAGGATG AAGCTCAAGAA |
| | Am *p53* forward | GGACTCTGATTAAGGCTCTGT |
| | Am *p53* reverse | ACCAGCAAAGGCAGAGGATT |
| | Am *p53* standard oligomer | GCAGGACTCTGATTAAGGCTCTGTCTGACCTA GGGCTGCAATGGTGCCTGTATTCGGTGCAATC CTCTGCCTTTGCTGGTGCT |
| | Am *agr* forward | TGTGAGTGAACAGTGCCAAGATG |
| | Am *agr* reverse | CTTGCTGGTTGTGAAATCTCGCA |
| | Am *agr* standard oligomer | ATTGTGAGTGAACAGTGCCAAGATGTCTGACC TCCTCCACCTCCTGCTTAGTTGCGAGATTTCAC AACCAGCAAGAA |
| | Nm *agr* forward | CAACATAATCCTGCTCTGTCGTGG |
| | Nm *agr* forward | GTTCCCGAAGTAGAAATCAAGTGG |
| | Nm *agr* standard oligomer | GACCAACATAATCCTGCTCTGTCGTGGGACTT GGCACGCCGGACGATGAATCCACTTGATTTCT ACTTCGGGAACATG |
| | Hd *agr* forward | CGCCCAGGAAGATGACTACAGTTG |
| | Hd *agr* forward | TGTTTTTACAGCCGTGAGGACC |
| | Hd *agr* standard oligomer | GCACGCCCAGGAAGATGACTACAGTTGATTTG ACATTCTCAAACCTGGTGTACATGCATAATTAA TATTATGGTCCTCACGGCTGTAAAAACACTT |
| | Amac *agr* forward | TGTGAGTGAACAGTGCCAAGATG |
| | Amac *agr* forward | TGGGAAGTCTCGCAACTAAGCAG |
| | Amac *agr* standard oligomer | ACTGTGAGTGAACAGTGCCAAGATGTCCGACC TCCTCCACCTCCTGCTTAGTTGCGAGACTTCCC AAA |
| *ORF primers* | Pw tORF forward$^{\#}$ | GTGCTCTCTGGGGGTTATAG |
| | Pw tORF reverse$^{\#}$ | AAAGTAGGTGTAAAGTGCACC |
| | Pw *cdkn2b* forward* $^{\#}$ | CATGTGAGGACACTGTTG |
| | Pw *cdkn2b* reverse* $^{\#}$ | TACTTCTGGATGGAGGAG |
| | Am tORF forward | CATGTCTCTGTGATAGC |
| | Am tORF reverse | CCCACGCATTAGAAGTAA |
| | Am *prod1* forward* | TGTAGTCTTTACAGGGTGACAT |
| | Am *prod1* reverse* | CACAAGCCTGCTACTTCTAGA |
